# Neuroimaging Biomarkers of Neuroprotection: Impact of Voluntary versus Enforced Exercise in Alzheimer’s Disease Models

**DOI:** 10.1101/2025.03.28.646015

**Authors:** Alexandra Badea, Ali Mahzarnia, Divya Reddy, Zijian Dong, Robert J Anderson, Hae Sol Moon, Jacques A Stout, Janai Williams, Lydianne Hirschler, Emmanuel L. Barbier, Christina L Williams

## Abstract

Exercise is a promising strategy for preventing or delaying Alzheimer’s disease (AD), yet its mechanisms remain unclear. We investigated how exercise influences brain structure, function, and behavior in a familial AD model. Mice underwent voluntary, voluntary plus enforced exercise, or remained sedentary. Neuroimaging included in vivo manganese-enhanced MRI (MEMRI). perfusion, and ex vivo diffusion MRI to assess morphometry, activity, cerebral blood flow (CBF), microstructural integrity and connectivity.

Both exercise regimens induced structural and functional brain adaptations while reducing anhedonia. Voluntary exercise increased cortical and limbic volumes, particularly in the hippocampus, cingulate, and entorhinal cortex, supporting cognitive and emotional regulation. Adding enforced exercise influenced subcortical and sensory regions, including visual, motor and associative areas, supporting sensory-motor integration. MEMRI revealed increased activity in sensorimotor, limbic, and associative cortices, with voluntary exercise enhancing limbic and associative regions, and enforced exercise strengthening sensorimotor and subcortical circuits.

White matter integrity improved in memory-associate pathways such as the corpus callosum, cingulum, and hippocampal commissure. Synaptic remodeling was observed in the cingulate cortex, anterior thalamic nuclei, and amygdala. Voluntary exercise enhanced CBF in the motor cortex and hippocampus, while enforced exercise limited these increases.

Connectivity analyses revealed exercise-responsive networks spanning the cingulate cortex, entorhinal cortex, anterior thalamic nuclei, and basolateral amygdala, and associated tracts. Graph analyses linked running distance with increased thalamic, brainstem, and cerebellar connectivity, associating exercise intensity with plasticity.

These findings highlight the ability of chronic exercise to modulate neuroimaging biomarkers through distinct but complementary pathways, reinforcing its potential as a neuroprotective intervention for AD.

**Highlights:**

- Exercise alters MRI biomarkers via distinct and partially overlapping mechanisms.
- Voluntary exercise boosts cortical and limbic regions for emotion and cognition.
- Enforced exercise strengthens subcortical and sensory areas for motor control.
- FA increases suggest memory tract reinforcement and grey matter remodelling.
- Graph analysis reveals plasticity in memory, emotion, and reward circuits.

## Introduction

Alzheimer’s disease (AD) is a complex, multifactorial condition with no effective treatments, driving increasing interest in strategies to reduce risk, prevent onset, and delay progression. AD prevalence was estimated to 6.9 million in the US, and this number is projected to double by 2050(Association 2024). AD is currently defined through biomarkers for proteinopathies leading to extracellular amyloid deposition and intracellular hyperphosphorylated tau, progressive neurodegeneration; as well as memory and behavioral impairments. AD is primarily characterized by learning and memory loss but is often accompanied by sensory-motor decline and behavioral changes, such as depression, social withdrawal, mood swings, and delusions, which may further increase risk. One common behavioral symptom is anhedonia (Lopez et al. 2003), a reduced ability to experience pleasure, which often accompanies depression. Anhedonia is increasingly recognized not only as a comorbidity but also as an early behavioral marker that may precede cognitive decline, and is accompanied by structural brain changes, such as cortical thinning (Clement, Wiborg, and Asuni 2020). Moreover, the rate of anhedonia increases from >36% in mild to moderate AD to 80% in severe AD (Saz et al. 2009).

While there has been limited success in slowing down cognitive decline and alleviating neuropsychiatric symptoms, there is robust evidence that exercise mitigates anhedonia and decreases inflammation, which affects neuronal signaling, reward decision making and other cognitive processes (Hird et al. 2024). Several other processes are associated with exercise and could influence cognition, for example increases in BDNF(Szuhany, Bugatti, and Otto 2015), synaptic plasticity (Lourenco et al. 2019; Vaynman, Ying, and Gomez-Pinilla 2004), altered neuroendocrine responses (Alghadir and Gabr 2020), decreased oxidative stress(Schuch et al. 2014) and neurovascular risk (Prins and Scheltens 2015), increase matter volume(Gujral et al. 2019) and myelination (Boa Sorte Silva et al. 2025). Moreover, animal models have shown effects on AD hallmark pathology (Adlard et al. 2005) (Derafshpour et al. 2025). There is therefore substantial interest in exploring non pharmaceutical preventive strategies that could reduce risk for AD in normal subjects at asymptomatic or MCI stages (Biazus-Sehn et al. 2020), understanding the mechanisms that lead to neuroprotection, and optimizing the exercise strategies that improve association to specific cognitive domains relevant to AD (Iso-Markku et al. 2024) (Chen, Zhao, and Zhou 2025; Raffin et al. 2025; Walker et al. 2025; Wang et al. 2025). It is unclear what are the causal relationships between exercise, biological brain changes and cognition, and what are optimal exercise strategies to maximize the effects of exercise on AD cognitive symptoms (Archer, Josefsson, and Lindwall 2014), and vulnerable brain circuits (David et al. 2025).

The hippocampus (Vaynman, Ying, and Gomez-Pinilla 2004) (Morris et al. 2017) (Bugg and Head 2011) (Derafshpour et al. 2025), has been at the core of AD studies, as it is a prime candidate and vulnerable brain region. One landmark cross species study in humans and mice showed increased cerebral blood volume and neurogenesis, in particular in the dentate gyrus (Pereira et al. 2007), in association with aerobic fitness and learning. Studies assessing the effects of exercise on AD risk could benefit from multimodal and network-based approaches to enhance sensitivity to subtle effects and uncover the governing principles of interactions between brain regions that work in synergy to sustain cognitive function (Burdette et al. 2010).

Despite strong evidence supporting exercise in mitigating AD pathology and symptoms, significant gaps remain in understanding how different types and intensities influence neurodegeneration. Most studies focus on broad interventions, with limited exploration of specific regimens and their effects on neural structure and function. Key neuroimaging markers—such as cerebral blood flow, connectivity, and microstructural integrity—are underexplored, particularly in AD models. Behavioral improvements, including cognitive resilience and mood regulation, show inconsistent findings, highlighting the need for systematic investigation. Notably, the impact of exercise on anhedonia and memory impairment—two core neuropsychiatric features of AD—remains understudied. Preliminary evidence suggests exercise may alleviate anhedonia via reward-related neural circuits and inflammatory pathways, yet mechanisms remain unclear. This study addresses these gaps through multimodal neuroimaging and behavioral assays to assess how varying exercise intensities shape brain structure, connectivity, and behavior in a familial AD model.

We utilized the CVN-AD mouse model (Colton et al. 2008b) (APPSweDI/mNos2^-/-^), which exhibits progressive cognitive deficits, neuroinflammation, amyloid and tau pathology (Wilcock et al. 2008), as well as hippocampal volume, and white matter microstructure (Badea et al. 2016) (Badea et al. 2019a) relative to controls. This model was chosen for its ability to mimic key aspects of AD progression, particularly the interplay between vascular dysfunction and neurodegeneration.

To assess the effects of exercise, we implemented two paradigms: voluntary wheel running and a combination of voluntary running with enforced voluntary+enforced exercise. The voluntary group engaged in daily wheel running, reflecting an active lifestyle, whereas the enforced exercise group underwent additional structured treadmill training, simulating high-intensity exercise. This design allowed us to compare the differential effects of exercise intensity and type on brain structure, connectivity, cerebral blood flow, and behavior. By integrating behavioral assessments with multimodal MRI, we aimed to elucidate the mechanisms through which exercise modulates AD-related changes and identify optimal strategies for mitigating disease progression.

Our primary objective was to determine how exercise influences neuroimaging biomarkers—cerebral blood flow, regional brain volume, and microstructural integrity—as well as behavioral outcomes such as memory and anhedonia. We hypothesized that enforced exercise would yield greater improvements in cognitive and neuropsychiatric symptoms compared to voluntary exercise alone. Additionally, we anticipated that exercise would enhance cerebral blood flow, structural integrity, and connectivity in regions associated with cognitive resilience. Given the limited data on exercise’s effects on brain connectivity in AD, we investigated whether different exercise intensities exert distinct effects on AD-related impairments.

By identifying how exercise modulates neurobiological markers and behavior, this study contributes to growing evidence supporting the role of physical activity in preserving cognitive function and brain integrity. Understanding the underlying neurobiological mechanisms can inform personalized lifestyle recommendations and therapeutic strategies aimed at reducing AD risk and slowing cognitive decline. Furthermore, our integration of neuroimaging and behavioral assessments provides a comprehensive framework for evaluating exercise-based interventions, potentially guiding clinical practices and public health policies that promote physical activity as a viable approach for AD prevention and management

## Methods

This study employed a multimodal imaging approach to investigate the effects of exercise on atrophy, microstructural integrity, and metabolism in CVN-AD mice. Structural MRI quantified regional brain volumes, diffusion MRI assessed microstructural integrity and connectivity, manganese-enhanced MRI measured neuronal activity, and perfusion MRI evaluated cerebral blood flow (CBF) as a proxy for metabolic function. Behavioral testing examined anhedonia and memory function. Female CVN-AD mice (APPSwDI/mNos2^-/-^), a model exhibiting progressive cognitive decline, amyloid and tau pathology, and neuroinflammation, were used in this study (Colton et al. 2008a; Wilcock et al. 2008; Colton et al. 2008b). The CVN-AD mice have mutated APP and a lowered expression of NOS2) and were created by Dr. Carol Colton at Duke University by crossing APPSweDI mice with mNos2^-/-^ mice to reduce immune differences between mice and humans, which are a worthwhile consideration in developing effective mouse models of AD, since immune-mediated levels of NOS2 mRNA and protein as well as levels of Nitric Oxide (NO) are significantly reduced in humans compared to mice. CVN-AD mice have mutated APP and a lowered expression of NOS2 to create a more human-like immune environment, to better mimic the age-dependent progression of AD pathology seen in humans. These pathological features include cerebral vascular amyloid deposition, tau pathology, and neuronal loss, as well as spatial memory deficits that develop over the first 52 weeks of life (Kan et al. 2015).

Mice were bred and housed at Duke University in a temperature-controlled facility with a reversed light-dark cycle (lights off at 8:00 AM) and had ad libitum access to standard chow and water. Mice were randomly assigned, by cages, to one of three experimental groups: sedentary (N=9), voluntary exercise (N=8), or voluntary+enforced exercise (N=9). Exercise training lasted 24 weeks, from 28 to 52 weeks of age. The voluntary exercise group had unrestricted access to running wheels in their cages for 6–8 hours per day, five days per week. The enforced exercise group followed the same protocol but also underwent treadmill training twice a week on non-consecutive days. The treadmill protocol consisted of a 5-minute warm-up (5 m/min), 35 minutes of running (10 m/min), and a 5-minute cool-down (5 m/min). Mice were not subjected to electric shocks, and motivation was provided by gentle prodding. The sedentary group remained in standard housing without exercise interventions. All procedures were approved by the Duke Institutional Animal Care and Use Committee (IACUC).

Behavioral assessments were conducted at 36 and 52 weeks of age to evaluate anhedonia and episodic memory. Anhedonia was assessed using the sucrose preference test, in which mice were given a choice between a water bottle and a 0.5% sucrose solution over four days. Bottle positions were alternated daily to prevent side bias, and sucrose preference was measured as the volume of sucrose solution consumed per day. Episodic memory was evaluated using the novel object recognition (NOR) test, which consisted of four trials: a habituation trial in an empty arena, two familiarization trials with two identical objects, and a novelty recognition trial where one familiar object was replaced with a novel object. Memory performance was quantified as the time spent exploring the novel object. To minimize social stress, mice were individually housed for one hour before testing, and all behavioral tests were conducted in a dimly lit, quiet room.

One week before imaging mice were implanted with Alzet 1007D minipumps (Durect Corp, Cupertino, CA), containing 100 μl of 64 μm/μl MnCl_2_*4(H_2_O) (Sigma Aldrich, St Louis, MO), in 100 mmol bicine (Sigma-Aldrich, St Louis, MO)(Badea et al. 2019b), using a ketamine xylazine cocktail for anesthesia (100:10 mg/Kg). Brain specimens were next prepared for ex vivo imaging. To reduce the T1 relaxation rate to ∼ 100 ms, we infused the brain with gadolinium via a transcardiac perfusion, followed by overnight formalin fixation, as in (Winter et al. 2024). Brain specimens were left in the skull to preserve brain shape, and avoid brain tissue damage, then imaged in fobmblin (Solvay).

In vivo MRI was performed using a 7.1T Bruker system (20 cm bore, 440 mT/m gradients, with AVIII console, and Paravision 6.0.1), with a volume excite and a 4 channel surface receive array coil. Mice were anesthetized with isoflurane (1–2%) and imaged using structural, diffusion, and perfusion MRI. Body temperature was maintained with warm circulating water. Structural MRI was acquired using a manganese enhanced (MEMRI) T1-weighted RARE sequence with TR = 300 ms, TE = 24.8 ms, RARE factor 4, and an isotropic resolution of 100 µm, with acquisition time of 22 minutes. Cerebral blood flow (CBF) was measured using pseudo-continuous arterial spin labeling (pCASL) as described in (Hirschler, Debacker, et al. 2018; Hirschler, Munting, et al. 2018),, with 30 pairs of control/label acquisitions (TR = 4500 ms, TE = 16.43 ms, labeling duration = 3000 ms, post-labeling delay = 300 ms). Fifteen axial slices (0.5 mm thick, 0.5 mm spacing) were acquired over two interleaved datasets in a total scan time of 9 minutes. Label and control phases were optimized via separate prescans. Inversion efficiency was estimated using FAIR-EPI in a single slice (0.25 mm in-plane resolution, 25 × 25 mm² field of view, 0.5 mm slice thickness) with the labeling plane placed 10 mm caudal to isocenter; the scan lasted 1 min 44 s. T1 values were estimated in representative animals using a 3D TrueFISP sequence (TR = 8 ms, TE = 4 ms), with 6 interleaved segments and a total scan repetition time of 2662 ms per inversion cycle. A quasi-geometric progression of 40 inversion times ranging from 119 to 2615 ms was used ([119, 126, 134, 141, 149, 157, 166, 175, 185, 195, 206, 217, 229, 241, 254, 268, 282, 297, 313, 330, 347, 365, 384, 404, 425, 447, 470, 494, 519, 545, 572, 600, 629, 659, 690, 722, 755, 789, 824, 861, 899, 938, 978, 1019, 1061, 1104, 1148, 1193, 1239, 1286, 1334, 1383, 1433, 1484, 1536, 1589, 1643, 1698, 1754, 1811, 1869, 1928, 1988, 2049, 2111, 2174, 2238, 2303, 2369, 2436,2504, 2573, 2615]). The field of view was 25 × 12.5 × 15 mm³, with a matrix of 100 × 50 × 30, yielding a resolution of 250 × 250 × 500 µm³. The total scan time was approximately 8 minutes. T1 maps were generated from signal recovery curves using a three-parameter exponential fit across the 40 TIs. A flow-compensated fast low-angle shot was used to estimate the inversion efficiency α, which was ∼0.75, by acquiring signal 3 mm rostral to the labeling plane. The fc-FLASH readout had TR/TE = 225/3.13 ms, with 2 averages and a total scan time of 1 minute and 44 seconds. The imaging geometry consisted of a single axial slice (1 mm thick) with an in-plane resolution of 234 × 234 µm² (matrix: 128 × 128, FOV: 30 × 30 mm²). To compute CBF, we applied the Buxton two-compartment model, using field-specific parameters (T1 of blood = 1.2 s at 7T, T1 of tissue = 0.6 s, λ = 0.9 mL/g). Control images from the ASL series were used to approximate fully relaxed tissue magnetization. As manganese shortens T1 and may bias CBF estimates, our analyses focused on relative CBF.

Fir ex vivo MRI we used a 9.4T (8.9 cm bore, 2000 mT/m gradients, with Agilent console), and in house built silver solenoid coils. We used a compressed sensing diffusion imaging protocol with TR/TE: 100 ms /14.2 ms; matrix: 420 x 256 x 256; FOV: 18.9 x 11.5 x 11.5 (mm), reconstructed at 45 um isotropic resolution, bandwidth: 62.5 kHz. We used 30 diffusion directions with b=4000 s/mm2; and 1 non diffusion weighted acquisition (b0). The max diffusion pulse amplitude was 130.57 G/cm; duration 4 ms; separation 6 ms, 8-fold compressed-sensing acceleration. The diffusion data was reconstructed (Anderson et al. 2018) (Wang et al. 2018) using the Berkeley Advanced Reconstruction toolbox (Tamir et al. 2016).

We used SAMBA for brain segmentation (Anderson et al. 2019), and a symmetrized ex vivo MRI brain atlas as a reference (Winter et al. 2024) https://zenodo.org/records/10652239. Regional volume analyses report volumes as % of the total brain volume. Voxel-based analyses were performed using SPM (Friston et al. 1994) and results were corrected for multiple comparisons using a cluster based false discovery rate (FDR) correction threshold of 0.05, before visualizing t or F maps. Tractography was done using DIPY with Q-ball Constant Solid Angle Reconstruction (Garyfallidis et al. 2014), the total brain was used as a mask, to assess connectivity. We used BRAINCONN2 (https://github.com/Ali-Mahzarnia/brainconn2) and DSI Studio (Yeh et al. 2013) for visualization.

All statistical analyses were performed using R (v4.2). Group differences were assessed using linear mixed models when tests were repeated, and linear models followed by ANOVA, and multiplicity correction using a false discovery rate threshold of 5%. To assess the effects of exercise and aging on sucrose preference, we conducted a linear mixed-effects model with treatment condition (sedentary, voluntary, and voluntary + enforced) and age (36 vs. 52 weeks) as fixed effects, and animal ID as a random effect. Linear models were use for single time point measurements, including image based regional metrics. FDR corrections were used to correct for multiplicity, and post hoc tests to compare group pairs after ANOVA. We focused on regions where the mean value in the voluntary wheel-running group was greater than in the sedentary group, to show beneficial effects of voluntary exercise. To prioritize the most affected brain regions, we ranked these regions based on significance, and the sum of the mean values across all three conditions (sedentary, voluntary, and voluntary+enforced exercise). This approach ensured that we highlighted statistically significant regions that were not only responsive to exercise but also demonstrated the strongest overall effects across conditions.

Connectivity differences were examined using sparse logistic regression (Wang, Zhang, and Dunson 2019) to compare groups (e.g., exercise vs. sedentary, enforced vs. voluntary exercise) and symmetric bilinear regression to evaluate the relationship between connectomes and running distance, as a continuous variable. Our approaches revealed subnetworks associated with exercise type as a categorical variable, and with exercise intensity, estimated by the running distance as a continuous variable.

## Results

### Body mass (Supplementary Figure 1)

High body mass and obesity are risk factors for AD, and our study examined how long-term exercise mitigates age related weight gain. The analysis across different exercise interventions and time points revealed significant effects of both treatment (F(2, 155) = 3.86, p = 0.023) and time (F(1, 140) = 10.18, p = 0.0017), along with a significant interaction between the two factors (F(2, 139) = 8.81, p = 0.0002). Sedentary mice exhibited significant weight gain from week 36 to week 52 (+1.97g, p < 0.00001, d = -5.37), whereas those in the voluntary + enforced exercise group maintained a lower body mass, weighing significantly less than sedentary mice at week 52 (-2.57g, p = 0.029, d = 2.96). At week 36, this exercise group also had lower body mass compared to sedentary mice at week 52 (-2.69g, p = 0.029, d = -3.10), suggesting a protective effect of exercise against age-related weight gain. Descriptive statistics further support these findings, with sedentary mice increasing from 23.84 ± 2.26g at week 36 to 25.59 ± 1.82g at week 52, while the voluntary + enforced group remained stable (22.90 ± 1.86g at Week 36 and 23.02 ± 1.38g at Week 52). Overall, these results highlight that long-term exercise mitigates age-related increases in body mass in mice.

### Anhedonia Reduction and Memory (Figure 1A, Supplementary Table 1)

Anhedonia, a hallmark neuropsychiatric symptom of AD, was significantly influenced by exercise. A linear mixed-effects model analysis revealed significant main effects of treatment (F(2, 218.5) = 22.76, p < 0.0001) and time (F(1, 218.5) = 34.15, p < 0.0001), with a treatment × time interaction (F(2, 218.5) = 9.83, p < 0.0001) suggesting that the anhedonia reduction was modulated by exercise. At 36 weeks, sedentary mice had a sucrose preference of 1.67 ± 1.16%, comparable to the voluntary exercise group (2.05 ± 1.21 %, t(91.99) = -1.08, p = 0.32). However, mice in the voluntary + enforced group exhibited a greater sucrose preference than the sedentary group (2.51 ± 1.16 %, t(91.99) = -2.39, p = 0.03), indicating that enforced exercise more effectively mitigated anhedonia. By 52 weeks, these effects became more pronounced. While the sedentary group showed only a modest and non-significant increase in sucrose preference (2.16 ± 1.32 %, t(220.5) = -1.43, p = 0.21), voluntary exercise led to a substantial improvement (4.07 ± 2.07%, t(91.99) = -6.66, p < 0.0001). The largest reduction in anhedonia was observed in the voluntary+enforced group, where sucrose preference more than doubled to 5.67 ± 2.11 %, t(91.99) = -9.03, p < 0.0001), suggesting that higher exercise engagement resulted in stronger reward-seeking behavior.

**Figure 1.**
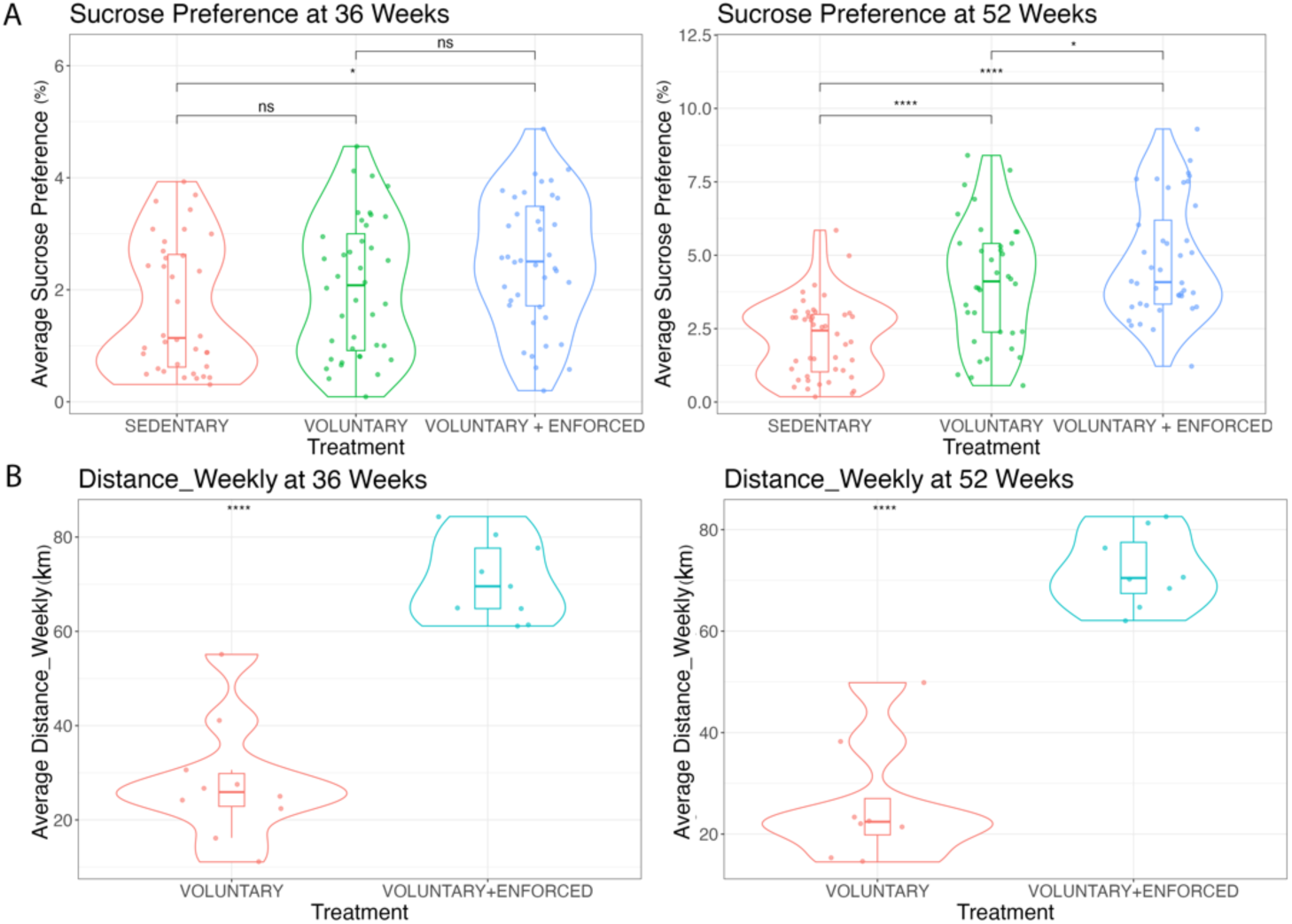
Effects of voluntary and enforced exercise on sucrose preference and exercise endurance. (A) Violin plots showing increasing sucrose preference between 36 and 52 weeks, and across three treatment groups: sedentary, voluntary, and voluntary+enforced. (B) Violin plots showing weekly running distance at 36 and 52 weeks for voluntary and voluntary + enforced groups. Mice in the voluntary + enforced condition exhibited greater weekly running distances than voluntary mice at both time points (p < 0.001). Boxplots within the violin plots indicate the median and interquartile range, with individual data measurement points overlaid. Statistical significance p values are FDR corrected: ns not significant, p < 0.05 (*), p<0.01 (**), p<0.001 (***).

Recognition indices (RI) in the Novel Object Recognition (NOR) test showed increases in the exercise groups compared to the sedentary group at both 36 and 52 weeks, with the largest difference observed in the voluntary+enforced group (+28.1% at 36 weeks, +37.0% at 52 weeks). However, these differences were not statistically significant (ns), indicating high variability within groups.

### Exercise Endurance and Training Adaptations (Figure 1B)

The reduction in anhedonia mirrored differences in exercise endurance, particularly in the voluntary+enforced exercise group. A linear mixed-effects model examined the effects of treatment (voluntary vs. voluntary+enforced) and time (36 vs. 52 weeks) on running distances, measured both daily and weekly. For daily running distance, neither the main effect of treatment (F(1,29.10) = 1.90, p = 0.179) nor time (F(1,14.13) = 1.49, p = 0.242) were significant. A marginal treatment × time interaction (F(1,14.13) = 4.11, p = 0.062) suggested a potential cumulative effect of enforced exercise over time. Weekly running distance revealed a strong treatment effect (F(1,30.40) = 40.89, p < 0.0001), indicating that mice in the voluntary + enforced group covered greater distances than the voluntary group. At 36 weeks, voluntary mice ran 28.00 ± 12.44 km/week, while voluntary + enforced mice ran more than twice this distance (70.78 ± 8.53 km/week). This difference remained at 52 weeks, with voluntary mice running 25.93 ± 12.05 km/week, whereas the voluntary + enforced mice sustained a higher distance of 72.01 ± 7.41 km/week. Importantly, these greater running distances aligned with anhedonia reduction. Mice that engaged in more total exercise (voluntary+enforced group) displayed the highest sucrose preference, reinforcing the link between endurance-based exercise and improvements in reward-seeking behavior.

### Volumetry (Figure 2)

We observed significant volume increases in multiple brain regions following both voluntary and enforced exercise, underscoring robust neuroplasticity. The total brain volume increased for both regimens, with voluntary+enforced exercise leading to a 9.6% increase (p = 4.35 × 10⁻⁷, η² = 0.477) and voluntary exercise producing a 10.5% increase relative to sedentary controls. This suggests widespread structural adaptations due to exercise. Specific brain regions exhibited pronounced changes, including the piriform cortex, involved in olfactory processing and associative learning. The voluntary+enforced exercise increased in volume by 14.4% (p = 1.12 × 10⁻⁷, η² = 0.514), and voluntary exercise led to a 13.2% increase compared to sedentary controls. Similarly, the ventral secondary auditory cortex, critical for auditory processing and sensory integration, showed an 18.3% increase for voluntary+enforced exercise (p = 7.57 × 10⁻⁸, η² = 0.524) and a 19.0% increase with voluntary exercise, supporting that exercise enhances sensory-related cortical structures. Exercise-related changes were also evident in regions involved in visual processing and higher-order cognitive function. The primary visual cortex (monocular area) exhibited a 10.9% increase following voluntarty+enforced exercise (p = 0.0059, η² = 0.177) and an 8.1% increase with voluntary exercise, indicating that physical activity may enhance visual processing circuits. Furthermore, the temporal association cortex, implicated in multisensory integration and high-level cognitive processing, showed a 10.9% increase in volume after voluntary+enforced exercise (p = 0.0026, η² = 0.205) and a 12.3% increase with voluntary exercise, suggesting that exercise may contribute to structural plasticity in regions linked to memory and executive function. The dorsolateral entorhinal cortex, linked to spatial memory and AD, demonstrated significant expansion, with voluntary+enforced exercise yielding a 19.8% increase (p = 1.63 × 10⁻⁵, η² = 0.370) and voluntary exercise producing a comparable 18.7% increase. This finding is relevant in the context of AD prevention, as the entorhinal cortex is critical for cognitive resilience. The significant effects in regions linked to memory, sensory, and higher cognitive processing highlight the potential of exercise as a non-pharmacological intervention for mitigating neurodegenerative decline.

**Figure 2.**
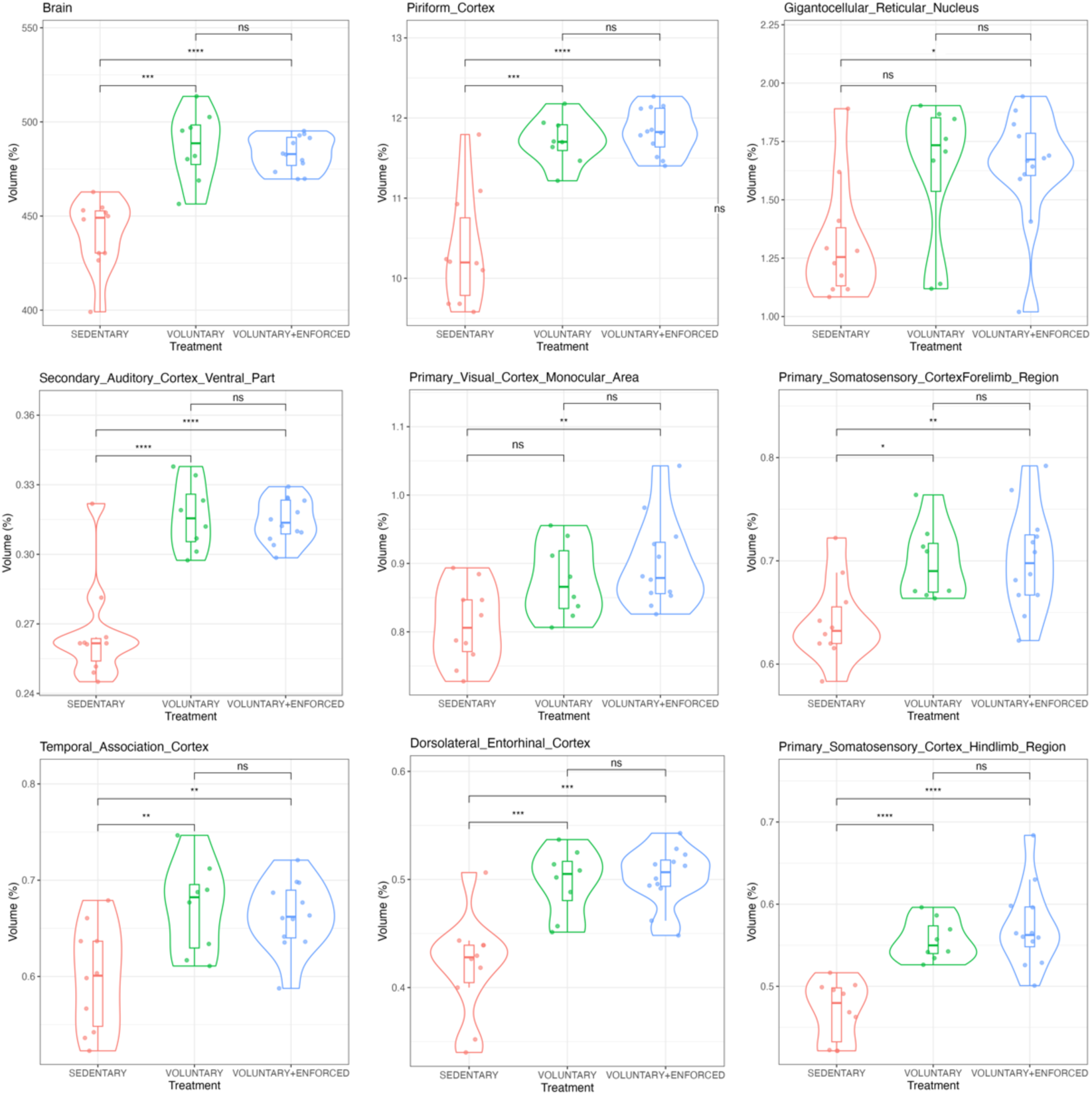
Exercise-induced regional volume increases. A. In vivo MRI revealed exercise induced significant increases in total brain volume, with major contributors including regions involved in sensory (piriform, V1, S1) and motor processing, autonomic and cardiovascular regulation (gigantocellular reticular nucleus), as well as memory and spatial navigation (entorhinal cortex). We also noted increases in the relative volumes of associative regions (entorhinal and temporal cortex). These findings highlight the structural plasticity associated with physical activity, particularly in regions involved in cognitive processing, perception, and motor coordination. Statistical significance p values are FDR corrected: ns not significant, p < 0.05 (*), p<0.01 (**), p<0.001 (***).

In addition, the primary somatosensory cortex (hindlimb region) also exhibited substantial plasticity, increasing in volume by 21.7% following voluntary+enforced exercise (p = 5.85 × 10⁻⁶, η² = 0.402) and 18.4% with voluntary exercise alone, reflecting the well-documented role of motor activity in shaping sensorimotor networks. In addition to these cortical regions, we observed exercise-induced changes in the gigantocellular reticular nucleus, a brainstem structure implicated in arousal and motor coordination. Voluntary+enforced exercise led to a 24.4% increase in volume (p = 0.0189, η² = 0.137), with a similar effect observed for voluntary exercise. This suggests that beyond higher-order cognitive regions, physical activity also impacts subcortical structures involved in movement regulation and autonomic function.

Certain subcortical structures exhibited relative volume reductions, highlighting region-specific adaptations to physical activity. Notably, the Cerebral Peduncle, involved in motor coordination and signal transmission between the brain and spinal cord, showed substantial volume decreases of 17.43% in voluntary+enforced-trained mice and 15.09% in voluntary exercise mice (F = 49.75, p < 0.0000001, η² = 0.632). These findings suggest that prolonged physical activity may induce plastic changes in white matter pathways, potentially reflecting increased signaling efficiency. Similarly, the Interpeduncular Nucleus, a midbrain structure involved in modulating reward and learning, displayed significant reductions of 33.27% (voluntary+enforced) and 31.31% (voluntary exercise) (F = 45.74, p < 0.0000002, η² = 0.612). Given its role in dopaminergic signaling and its connections with the habenula, this may indicate an adaptive response to altered metabolic demands. Volume reductions were also observed in the Ventral Tegmental Area (VTA), a critical region for motivation, reward processing, and reinforcement learning, with decreases of 30.28% in voluntary+enforced mice and 30.39% in voluntary exercise mice (F = 31.41, p < 0.0000024, η² = 0.520). Finally, the Dorsal Tegmentum, involved in autonomic regulation and motor coordination, exhibited 10.16% and 11.28% volume reductions in voluntary+enforced and voluntary exercise groups, respectively (F = 32.65, p < 0.0000024, η² = 0.530). These changes suggest that exercise selectively modulates brainstem circuits, possibly optimizing connectivity between motor and autonomic control centers. These findings indicate that while exercise enhances structural integrity in many regions, it also leads to relative volume decreases in select subcortical areas, particularly those involved in motor control, reward processing, and autonomic regulation. These reductions may represent structural refinements in neural networks, potentially improving functional efficiency and metabolic homeostasis in response to increased physical activity. **Supplementary Table 2** summarizes the mean regional volume values, standard deviations, effect sizes (Cohen’s F), and statistical comparisons.

Ex vivo analyses revealed exercise-induced volume increases in regions such as the pretectal nucleus and pre- and postsubiculum, suggesting neuroprotective effects (**Supplementary Table2, and Supplementary Figure 2**). The higher image resolution enabled the detection of significant changes in white matter tracts, including the anterior commissure and facial nerve (η² =0.3, p<0.01), which were not reliably detected in vivo. Voluntary exercise primarily benefited cortical structures, while enforced exercise had a more pronounced impact on subcortical and sensory regions. The pretectal nucleus exhibited the largest effect size (η² = 0.36, F(2, 81) = 16.99, p<0.005), with both voluntary and enforced exercise associated with increased volume compared to sedentary controls. Notably, the anterior commissure (η² = 0.28, F(2, 81) = 11.47, p<0.01) and the facial nerve (η² = 0.30, F(2, 81) = 13.10, p<0.01) also showed significant volumetric increases, underscoring white matter and cranial nerve plasticity with activity. Exercise-related expansion was further observed in the postsubiculum (η² = 0.31, F(2, 81) = 13.57, p<0.01), a hippocampal-adjacent region critical for spatial navigation, and in the presubiculum (η² = 0.21, F(2, 81) = 8.06, p<0.05), highlighting recruitment of memory-related circuitry. Subcortical sensory relay centers were also sensitive to activity level: the medial geniculate (η² = 0.25, F(2, 81) = 9.95, p<0.05), parabrachial (η² = 0.25, F(2, 81) = 9.89, p<0.01), and latero-posterior thalamic nuclei (η² = 0.21, F(2, 81) = 7.95, p<0.05) all demonstrated significant volume differences. Lastly, the microcellular tegmental nucleus, involved in autonomic regulation, showed increased volume with exercise (η² = 0.20, F(2, 81) = 7.70, p<0.05), suggesting broader engagement of brainstem structures. Post hoc analyses revealed that both voluntary and combined (voluntary + enforced) exercise groups exhibited significant increases compared to sedentary controls, with the combined exercise group showing the most robust effects. The largest enhancements were observed in the Postsubiculum, Pretectal Nucleus, and Anterior Commissure, with increases of up to 5.4% relative to sedentary mice (*p* < 0.0001), implicated in spatial memory, multisensory integration, and interhemispheric communication. These results suggest that exercise promotes structural plasticity in circuits relevant to cognition and sensorimotor processing. The combined exercise produced stronger effects than voluntary exercise in regions including the Pretectal and Parabrachial nuclei—involved in visual reflexes and autonomic regulation. A comparison between the two exercise groups showed a larger effect of the combined exercise in the Medial Geniculate Nucleus (*p* = 0.0063), a relay center for auditory information. These results suggest that adding structured exercise amplifies the impact across sensory, motor, and cognitive circuits.

Voxel based analysis results supported the regional volumetry findings (**Figure 3**), indicating widespread exercise effects. We noted increases across olfactory and orbital areas, the piriform, auditory and auditory cortices, M1 and S1, tenia tecta, septum, bed nucleus of stria terminalis, anterior thalamic nuclei, hypothalamus, parietal association cortex, hippocampus, amygdala and entorhinal and cingulate cortices, midbrain reticular nuclei, inferior colliculus, V1, pons, and cerebellum. Volume increases in exercised mice relative to sedentary indicated a role for olfactory areas, S1, M1, piriform cortex, insular and auditory cortex, ventral hippocampus, V1, cerebellum and pons. Voluntary exercised mice had larger volumes in the dorsal hippocampus, right M1 and S1, striatum, and accumbens, as well as superior colliculus, V1, subiculum, and cingulate cortex, inferior colliculus and cerebellum. Voluntary plus enforced mice had larger regional volumes on the right S1, bed nucleus of stria terminalis, temporal association area, medial geniculate, midbrain reticular nuclei.

**Figure 3.**
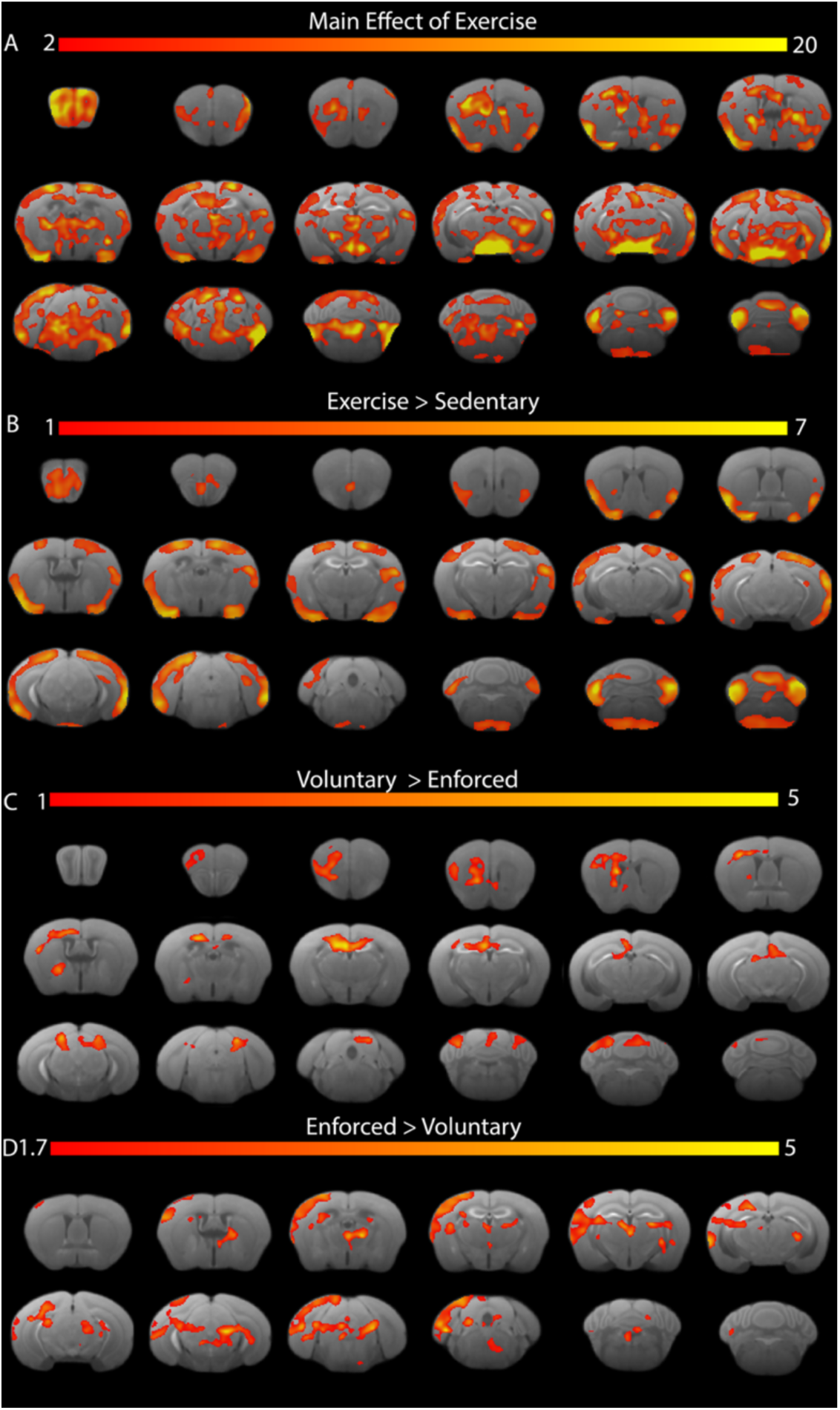
In vivo voxel based morphometry effects of exercise and exercise type. A. Main effect of exercise. B. Exercise > Sedentary. C. Voluntary > Voluntary +Enforced. D. Voluntary + Enforced > Voluntary. Statistical maps are thresholded at FDR=5%.

Ex vivo voxel-based analysis (VBA) refined the focus of effects observed in vivo, and confirmed volume increases in the dorsal hippocampus, showing layer specific effects, and the cerebellum in voluntary exercise mice. Additionally, it revealed effects in the ventral hippocampus, piriform cortex, and pyramidal tracts. The enforced + voluntary exercised mice showed further volumetric increases in the secondary visual cortex (V2) and more pronounced effects in the cingulate and primary motor (M1) cortices (**Supplementary Figure 3**). Mice subjected to enforced exercise had enlarged olfactory areas, M1/S2, and parasubiculum. Voluntary exercised mice had enlarged cingulate cortex, temporal, auditory, and piriform cortex, dorsal and ventral hippocampal areas, cerebellum, brain stem, and superior colliculus, relative to voluntary + enforced mice. Enforced plus voluntary exercised mice showed clusters in the olfactory areas, thalamus, amygdala and visual cortex.

Exercise-induced microstructural changes, and revealed significant fractional anisotropy (FA) increases in multiple sensorimotor regions, suggesting enhanced white matter integrity and connectivity (**Figure 4, Supplementary Table 3**). The most pronounced FA increases were observed in the primary somatosensory cortex, particularly in the jaw region (F = 5.06, η² =0.14, p = 0.048, +11.7% voluntary, -6.7% voluntary+enforced), upper lip region (F = 7.78, η²=0.21, p = 0.037, +10.2% voluntary, -8.4% voluntary+enforced), and barrel field (F = 5.95, η²=0.17, p = 0.03, +6.5% voluntary, -4.8% voluntary+enforced). The medial lemniscus, a critical relay for proprioceptive and somatosensory information, also exhibited an FA increase with voluntary exercise (F = 6.08, η²=0.17, p = 0.03, +5.2% voluntary), whereas enforced exercise was associated with a decrease (-7.3%). A similar pattern was noted in the secondary somatosensory cortex (F = 6.67, η²=0.18, p = 0.03, +3% voluntary, -14.7% voluntary+enforced) and secondary auditory cortex (ventral part) (F = 7.67, η² =0.20, p = 0.03, +1% voluntary, -11.2% voluntary+enforced), indicating that exercise-driven microstructural plasticity extended beyond sensorimotor pathways to regions involved in auditory processing. Marginally significant FA increases (η² ∼0.1, p < 0.1) were observed in the frontal cortex (area 3) (+10.93% voluntary, -9.1% voluntary+enforced), insular cortex (+5.33% voluntary, -11.2% voluntary+enforced), corpus callosum (+4.12% voluntary, -4.1% voluntary+enforced), and cingulum (+3.89% voluntary, -7.0% voluntary+enforced).

**Figure 4.**
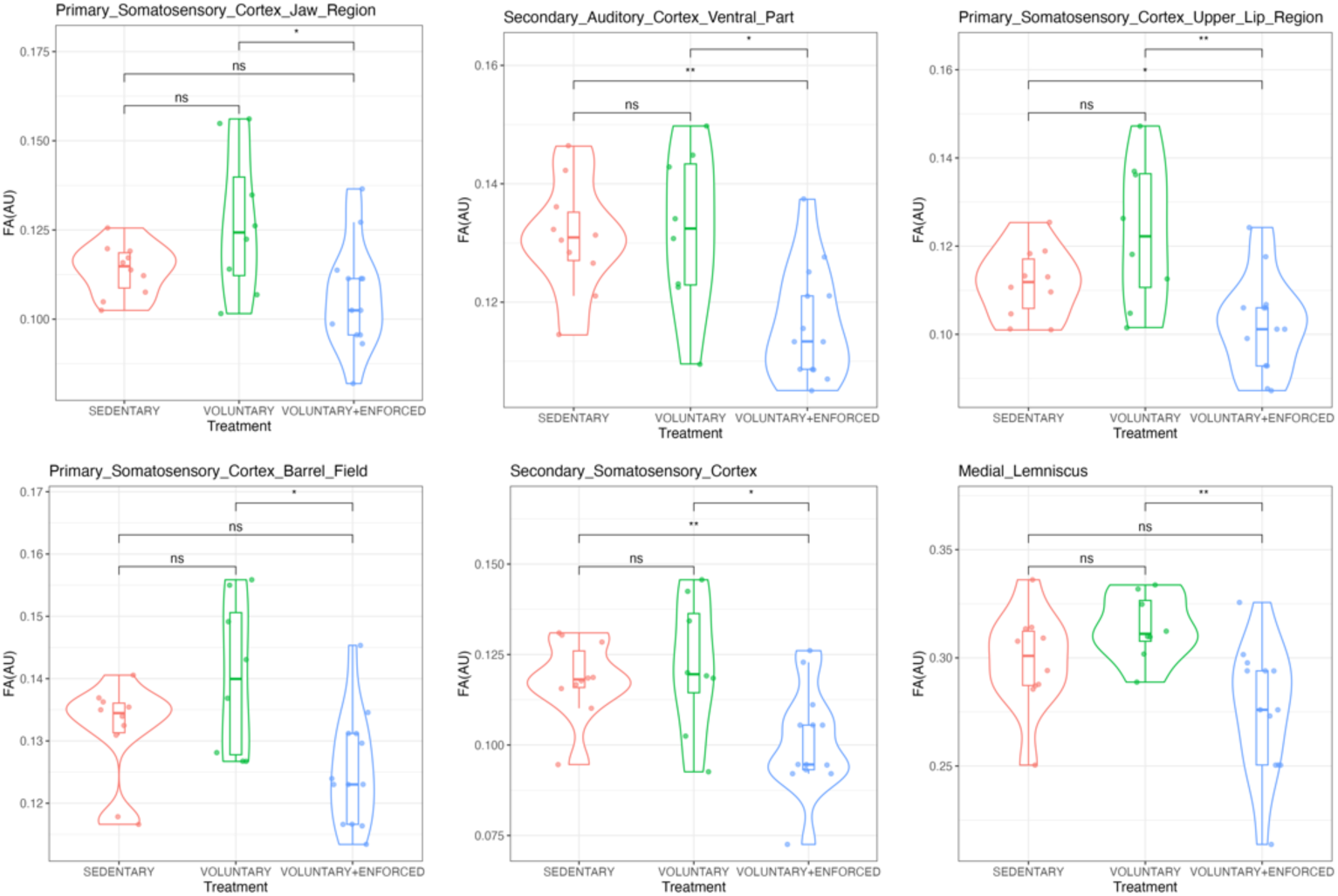
Fractional anisotropy (FA) increases across brain regions following different exercise regimens. FA values were highest in the voluntary exercise group compared to the sedentary condition in multiple sensory and motor regions, with notable differences in the Primary Somatosensory Cortex (Jaw and Upper Lip Regions), Medial Lemniscus, and Secondary Somatosensory Cortex. Boxplots within the violins indicate the interquartile range, median, and data distribution. Statistical significance is based on ANOVA followed by posthoc pairwise comparisons with false discovery rate (FDR) correction. Regions with significant differences (FDR-corrected p-values) are indicated with asterisks. Statistical significance: ns not significant, p < 0.05 (*), p<0.01 (**), p<0.001 (***).

Exercise-induced microstructural changes also revealed significant FA decreases across several subcortical and limbic regions, particularly when adding enforced exercise. The most pronounced FA reductions were observed in the laterodorsal nucleus of the thalamus (F = 11.21, η² = 0.27, FDR-corrected p = 0.0095, −15.1%) and the anterior thalamic nuclei (F = 11.14, η² = 0.27, FDR-corrected p = 0.0095, −14.8%), key relay structures involved in memory and attention. Additional thalamic and brainstem regions, including the pedunculotegmental and medial paralemniscial nuclei (F = 8.28, η² = 0.22, FDR-corrected p = 0.0257, −14.7%), cuneiform nucleus (F = 6.00, η² = 0.17, FDR-corrected p = 0.0326, −13.9%), and bed nucleus of the stria terminalis (F = 6.35, η² = 0.17, FDR-corrected p = 0.0326, −13.2%), also exhibited significant FA decreases. Limbic regions such as the cingulate cortex (area 25; F = 6.02, η² = 0.17, FDR-corrected p = 0.0326, −13.0%), ventral pallidum (F = 7.07, η² = 0.19, FDR-corrected p = 0.0323, −12.7%), and amygdala (F = 8.09, η² = 0.21, FDR-corrected p = 0.0259, −11.5%) were similarly affected. Notably, the primary somatosensory cortex (hindlimb region) also showed a significant FA reduction (F = 7.34, η² = 0.20, FDR-corrected p = 0.0297, −11.2%), suggesting that sensorimotor regions are not uniformly strengthened by physical activity and may be susceptible to the mode of exercise. Additional decreases in regions such as the claustrum, preoptic telencephalon, septum, ventral hippocampal commissure, and facial nerve further underscore the widespread impact of enforced exercise on brain microstructure. We note that voluntary exercise often mitigated the declines observed with the addition of enforced exercise, bringing FA levels closer to sedentary controls. These results underscore the need to differentiate between voluntary and enforced exercise effects on brain microstructure.

Post hoc analyses revealed significant reductions in FA in multiple brain regions in voluntary plus enforced exercised animals compared to sedentary controls, suggesting long-term microstructural adaptations. The most pronounced effects were detected in the Latero-Dorsal Nucleus of the Thalamus (t = 4.53, p = 0.0003), Hippocampus (t = 4.23, p = 0.0006), and Cingulate Cortex Area 24a (t = 4.19, p = 0.0007), crucial for sensory integration, memory, and executive function. Additional FA reductions were found in the Amygdala (t = 3.98, p = 0.0013), Substantia Nigra (t = 3.69, p = 0.0026), and Anterior Thalamic Nuclei (t = 3.68, p = 0.0027), suggesting remodeling of circuits involved in emotion regulation and motor control. Comparing voluntary exercise to sedentary conditions revealed significant FA reductions across multiple brain regions, such as the Anterior Thalamic Nuclei (t = 4.40, p = 0.0004), Facial Nerve (t = 4.15, p = 0.0008), and Cingulate Cortex Area 25 (t = 3.47, p = 0.0047). Additional FA decreases were detected in the Latero-Dorsal Nucleus of the Thalamus (t = 3.46, p = 0.0048), Bed Nucleus of the Stria Terminalis (t = 3.30, p = 0.0071), and Pedunculotegmental Nuclei (t = 3.11, p = 0.0116). These structures contribute to arousal regulation, reward processing, and stress adaptation, implying that voluntary exercise could drive reorganization in circuits associated with motivation and autonomic regulation. Notably, the Claustrum (t = 2.67, p = 0.0322) and Preoptic Telencephalon (t = 2.65, p = 0.0333), both involved in sensory integration and thermoregulation, also showed FA reductions following voluntary exercise.

The comparison between voluntary exercise alone and combined voluntary + enforced exercise revealed FA reductions in several sensorimotor and associative regions, suggesting that adding enforced exercise may modify or counteract some of the neuroplastic adaptations associated with voluntary activity. Notably, FA reductions were observed in the Primary Somatosensory Cortex (Upper Lip Region: t = 3.93, p = 0.0014; Jaw Region: t = 3.18, p = 0.0097) and the Barrel Field (t = 3.45, p = 0.0050), indicating that the microstructural integrity of key sensorimotor processing areas was more strongly enhanced by voluntary exercise alone compared to a combined paradigm. Similarly, the Medial Lemniscus (t = 3.39, p = 0.0057), responsible for relaying somatosensory and proprioceptive information, exhibited greater FA with voluntary exercise than when enforced exercise was introduced. FA reductions in the Secondary Somatosensory Cortex (t = 3.24, p = 0.0083) and Secondary Auditory Cortex (t = 3.31, p = 0.0070) further indicate that voluntary exercise alone may promote stronger microstructural adaptations in regions beyond primary motor pathways, with potential implications for sensory integration and higher-order processing. More modest FA reductions were detected in the Frontal Cortex (Area 3: t = 2.86, p = 0.0208), Ventral Thalamic Nuclei (t = 2.82, p = 0.0229), Corpus Callosum (t = 2.64, p = 0.0346), and Zona Incerta (t = 2.57, p = 0.0401). These regions are involved in cognitive control, interhemispheric communication, and arousal regulation, suggesting that enforced exercise may alter neuroplastic adaptations in both sensorimotor and cognitive circuits. These results support the idea that voluntary exercise alone induces pronounced microstructural adaptations, while enforced exercise may impose competing mechanisms of plasticity.

Voxel-based analysis (VBA) of fractional anisotropy (FA) revealed a significant effect of exercise in the olfactory areas, motor cortices, tenia tecta, insula, cingulate cortex, septum, hippocampus, amygdala, piriform cortex, primary visual cortex (V1), hypothalamus, substantia innominata, periaqueductal gray, cerebellum, pons, and pyramidal tract (**Figure 5**). Voluntary exercise was associated with greater FA increases in the corpus callosum, cingulum, fornix, zona incerta, subthalamic nuclei, hippocampus, hippocampal commissure, and cingulate cortex compared to voluntary+enforced conditions. Our findings highlight a strong overlap between volumetric increases and FA alterations, suggesting that complex mechanisms of exercise-induced neuroplasticity extend to both gray and white matter compartments.

**Figure 5.**
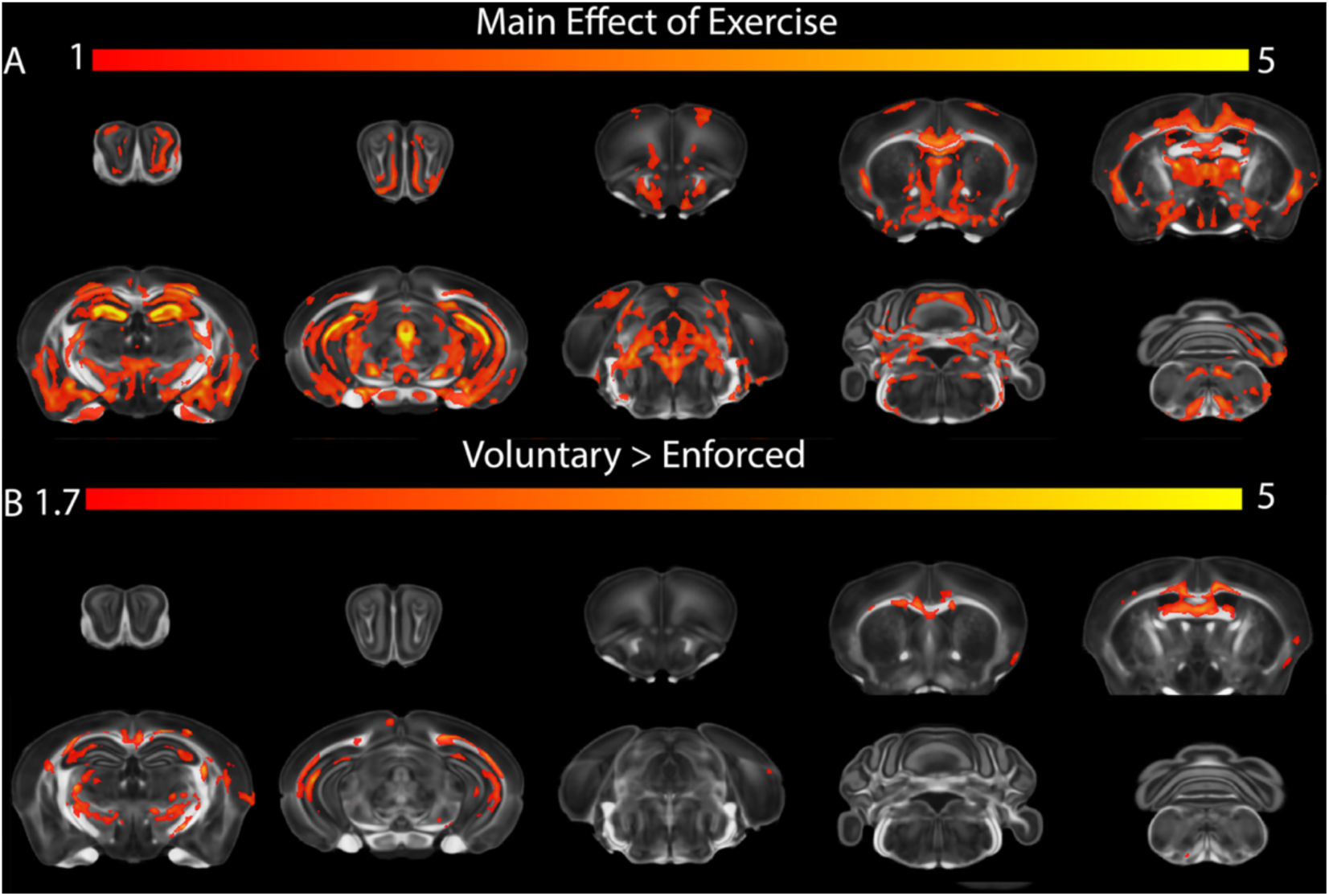
Effects of exercise on voxel based fractional anisotropy estimates. Exercise induced changes in olfactory, motor areas, cingulate cortex as well as septum, thalamus, accumbens, amygdala and cerebellum. A larger positive impact of exercise was observed in voluntary trained relative to voluntary + enforced trained mice in the corpus callosum, cingulum, fornix, zona incerta, hippocampus and cingulate cortex. FDR=5%.

Manganese-enhanced MRI (MEMRI) revealed widespread and region-specific alterations in T1w signal intensity in response to both voluntary wheel running and enforced treadmill exercise, highlighting distinct neuroplastic responses across the brain (**Figure 6, Supplementary Table 4**). Signal increases were most prominent in cortical regions involved in sensorimotor integration, cognition, and sensory processing, whereas decreases were more evident in brainstem and cerebellar structures, especially under enforced conditions. Frontal, orbital, insular, and somatosensory cortices exhibited the most robust increases in manganese uptake.

**Figure 6.**
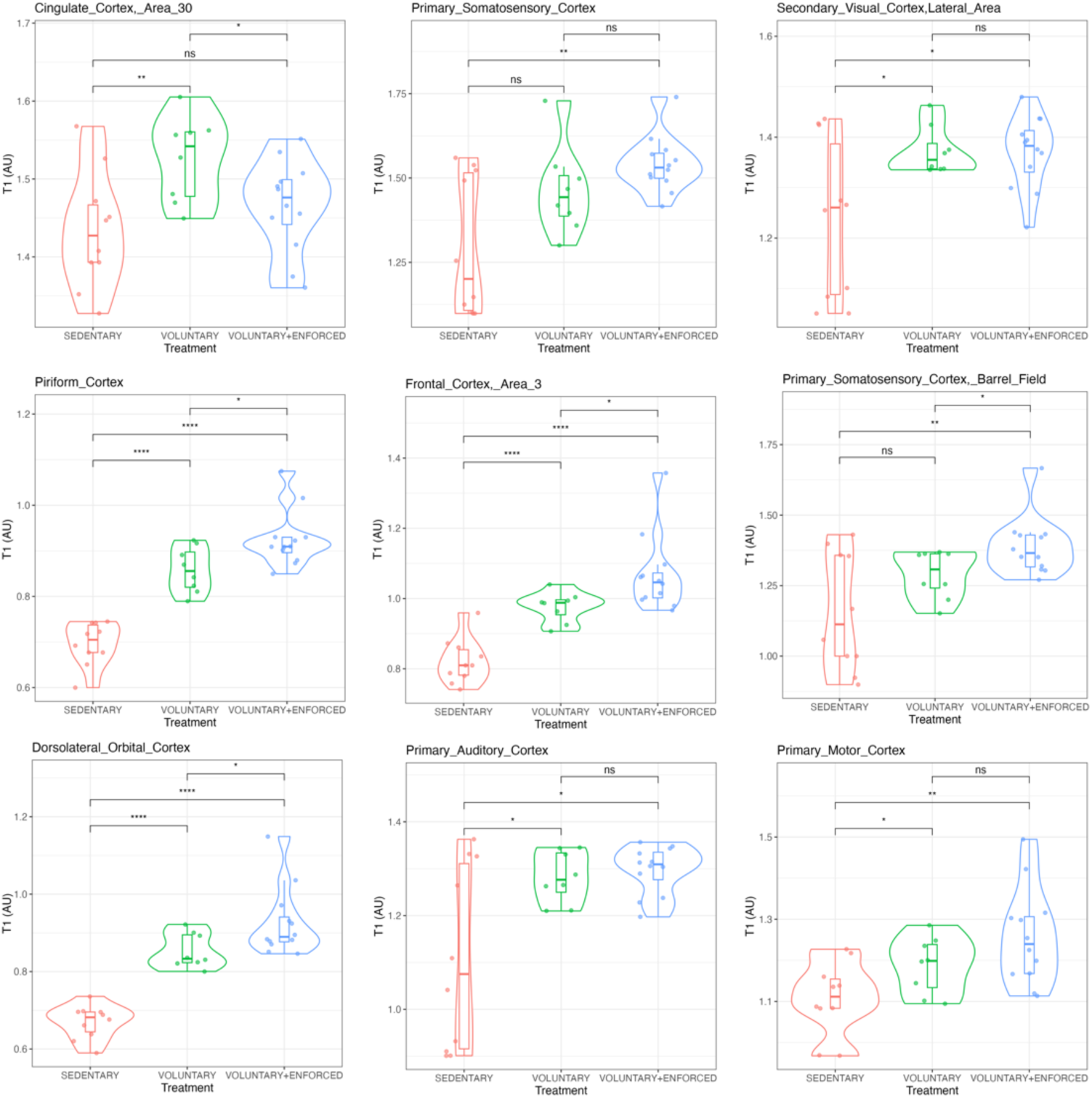
T1w MEMRI signal intensity across brain regions in exercise groups compared to the sedentary conditions. The largest differences were noted in somatosensory and motor areas, the cingulate cortex, as well as auditory cortices. Differences between groups are indicated by horizontal bars with asterisks Statistical significance p values are FDR corrected: ns not significant, p < 0.05 (*), p<0.01 (**), p<0.001 (***).

The piriform cortex showed the strongest post hoc difference between sedentary and exercised animals (p = 6.1×10⁻¹⁰), accompanied by a large overall effect (F = 49.65, η² = 0.63, p = 1.4×10⁻⁷; +32.8% voluntary+enforced, +23.2% voluntary). Similarly, the dorsolateral orbital cortex showed pronounced post hoc significance (p = 2.8×10⁻⁹) and large effect size (F = 42.98, η² = 0.60, p = 3.2×10⁻⁷; +38.3% voluntary+enforced, +27.4% voluntary). Significant post hoc effects were also observed in the lateral orbital cortex (p = 2.0×10⁻⁸), frontal association cortex (p = 2.2×10⁻⁸), and frontal cortex, area 3 (p = 3.9×10⁻⁷), all showing strong signal increases in both voluntary+enforced and voluntary conditions. The insular cortex, ventral claustrum, and lateral olfactory tract also showed increased signal, underscoring engagement of associative and integrative cortices. Within the somatosensory system, the primary somatosensory cortex (jaw region: F = 18.78, η² = 0.39, p = 1.1×10⁻⁴; +28.6% voluntary+enforced, +17.2% voluntary; upper lip: F = 11.16, η² = 0.28, p = 0.0013; +26.4% voluntary+enforced, +15.1% voluntary) was significantly affected, as were the dysgranular zone and forelimb region (F = 6.05 and 4.94, η² = 0.17 and 0.15; p = 0.0119 and 0.0245), reflecting widespread somatosensory plasticity. The primary motor cortex was also modulated (F = 6.45, η² = 0.18, p = 0.0096), particularly in response to combined exercise. Auditory and visual processing areas were similarly engaged. The secondary auditory cortex, both ventral (F = 9.38, η² = 0.24, p = 0.0022) and dorsal parts (F = 7.56, η² = 0.21, p = 0.0053), as well as the primary auditory cortex (F = 7.67, η² = 0.21, p = 0.0052), showed increased signal intensity with exercise. The secondary visual cortex (lateral area) also showed enhanced signal (F = 5.42, η² = 0.16, p = 0.0183; +10.7% voluntary+enforced, +11.0% voluntary), highlighting plasticity in sensory networks. Cognitive and associative brain regions also exhibited significant exercise-driven signal increases. The cingulate cortex demonstrated consistent effects across Area 32 (F = 9.65, η² = 0.25, p = 0.0019), Area 30 (F = 4.78, η² = 0.14, p = 0.0270), and Area 24a (F = 4.07, η² = 0.12, p = 0.0445), implicated in attention, executive function, and motivational salience. The hippocampus, critical for learning and memory, was also significantly affected (F = 7.44, η² = 0.20, p = 0.0055). Additional subcortical regions including the striatum (F = 16.04, η² = 0.36, p = 2.6×10⁻⁴), accumbens (F = 18.10, η² = 0.38, p = 0.0001), and claustrum (F = 23.69, η² = 0.45, p = 2.6×10⁻⁵) underscore the engagement of reward and integration circuits. Voluntary+enforced exercise was also associated with reductions in T1w signal in brainstem and cerebellar structures, suggesting downregulation of integrative and relay centers. The spinal trigeminal nucleus exhibited the strongest decrease (F = 10.89, η² = 0.27, p = 0.0013; −41.1% voluntary+enforced, −33.7% voluntary), followed by the spinocerebellar tract (F = 11.07, η² = 0.28, p = 0.0013; −36.9% voluntary+enforced, −36.6% voluntary), and cuneate nucleus (F = 10.27, η² = 0.26, p = 0.0016; −36.4% voluntary+enforced, −26.2% voluntary). Significant suppression was observed in the dentate (F = 9.89, η² = 0.25, p = 0.0018; −26.0% voluntary+enforced, −13.6% voluntary) and fastigial nuclei (F = 10.75, η² = 0.27, p = 0.0013; −30.1% voluntary+enforced, −17.8% voluntary). Posthoc comparisons indicated that enforced exercise may have distinct modulatory effects beyond those induced by voluntary activity alone for regions e.g. the Lateral Orbital Cortex, Claustrum, and Ventral Orbital Cortex. These findings support that different exercise paradigms modulate multiple networks—including sensory, motor, cognitive, and autonomic systems. Enforced exercise was associated with greater enhancement in cortical and limbic structures and suppression in subcortical and brainstem regions, suggesting a redistribution of neural activity toward higher-order integrative circuits.

Voxel-based analysed showed that exercise increased T1w signal in areas implicated in sensorimotor integration, emotional regulation, and autonomic control, including olfactory areas, motor cortex, accumbens, septum, preotpic area and hypothalamus, amygdala, as well as ventral hippocampus, subicullum, S1 and periaquductal gray, and trigeminal reticular nucleus (**Figure 7**). Voluntary mice had higher signal in the cingulate cortex, posterior parietal association area, superior coliculus, motor related areas, cerebellum and brain stem. Conversely, enforced exercise had larger effects localized to the left M1 and S1, corticoamygdalar area and brain stem. Results reflect divergent neuroplasticity patterns across exercise modalities.

**Figure 7.**
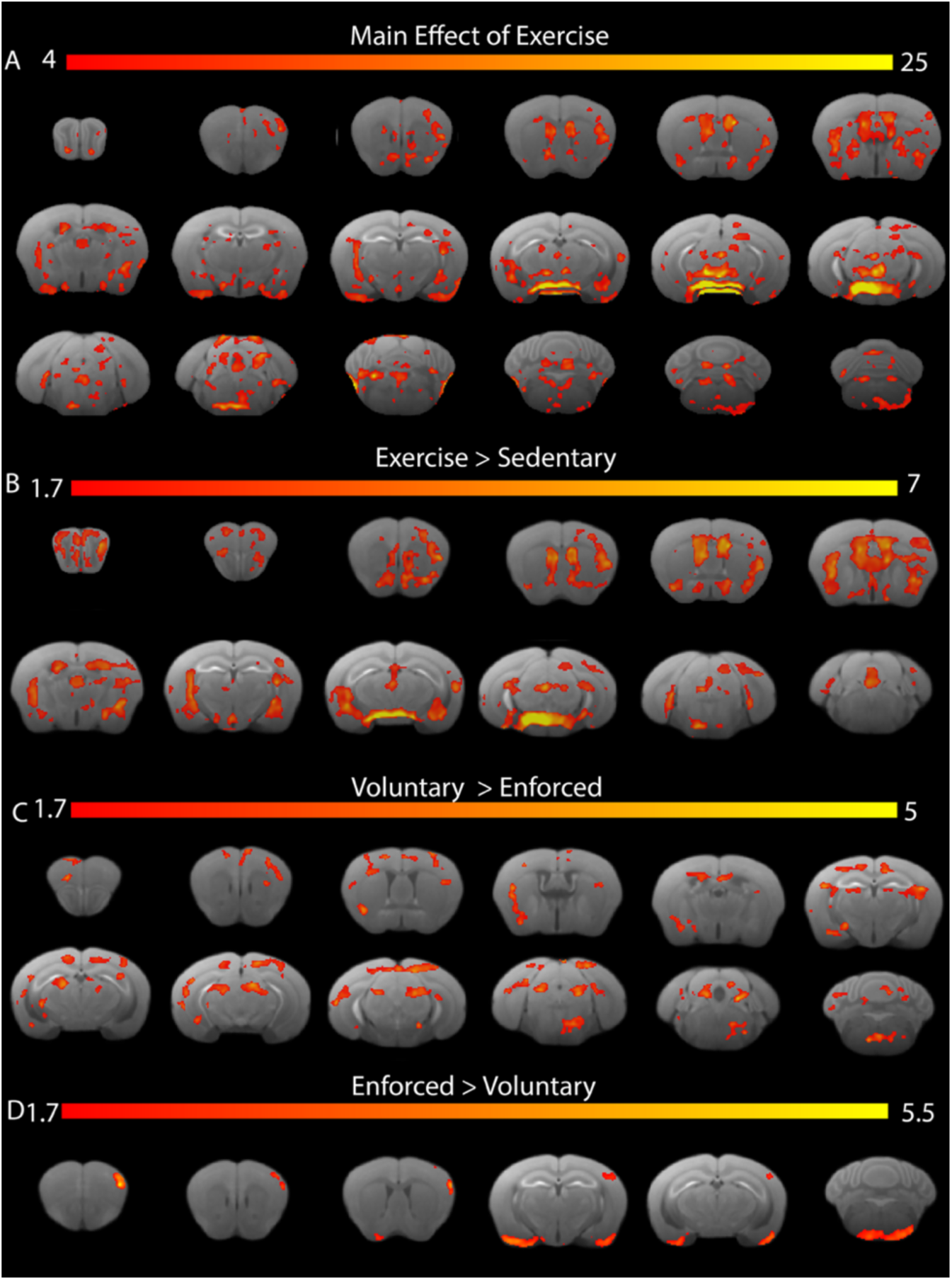
VBA analyses indicated that exercise impacted manganese uptake. Exercise impacted the olfactory areas, motor cortex, accumbens, septum, preotpic area and hypothalamus, amygdala, ventral hippocampus, subicullum, S1, periaquductal gray, and trigeminal reticular nucleus. Voluntary mice had higher signal in the cingulate cortex, posterior parietal association area, superior coliculus, motor related areas, cerebellum and brain stem. Enforced exercise induced increases in the left M1 and S1, corticoamygdalar area and brain stem. FDR=5%.

Regional analyses suggest that exercise may modulate subcortical perfusion, with marginal trends observed in the Ventral Thalamic Nuclei, Superior Colliculus, and Midbrain Reticular Nucleus (p=0.1), all functionally linked to motor control, sensorimotor integration, and locomotor arousal. Although not statistically significant after correction, the consistency in direction and effect size (η²∼0.2) supports a potential role for exercise in enhancing regional vascular response, in a way that may not be evenly distributed across brain regions.

In vivo VBA captured changes in relative cerebral blood flow (**Figure 8**), for extensive cortical domains, such olfactory areas, motor and visual cortices, tenia tecta, septum, striatum, hypothalamus, and temporal lobe structures including amygdala, hippocampus, and thalamus, as well as cerebellum and brain stem. Voluntary exercised mice had increased CBF in the right motor cortex, dorsal hippocampus, accumbens, visual cortex and inferior colliculus.

**Figure 8.**
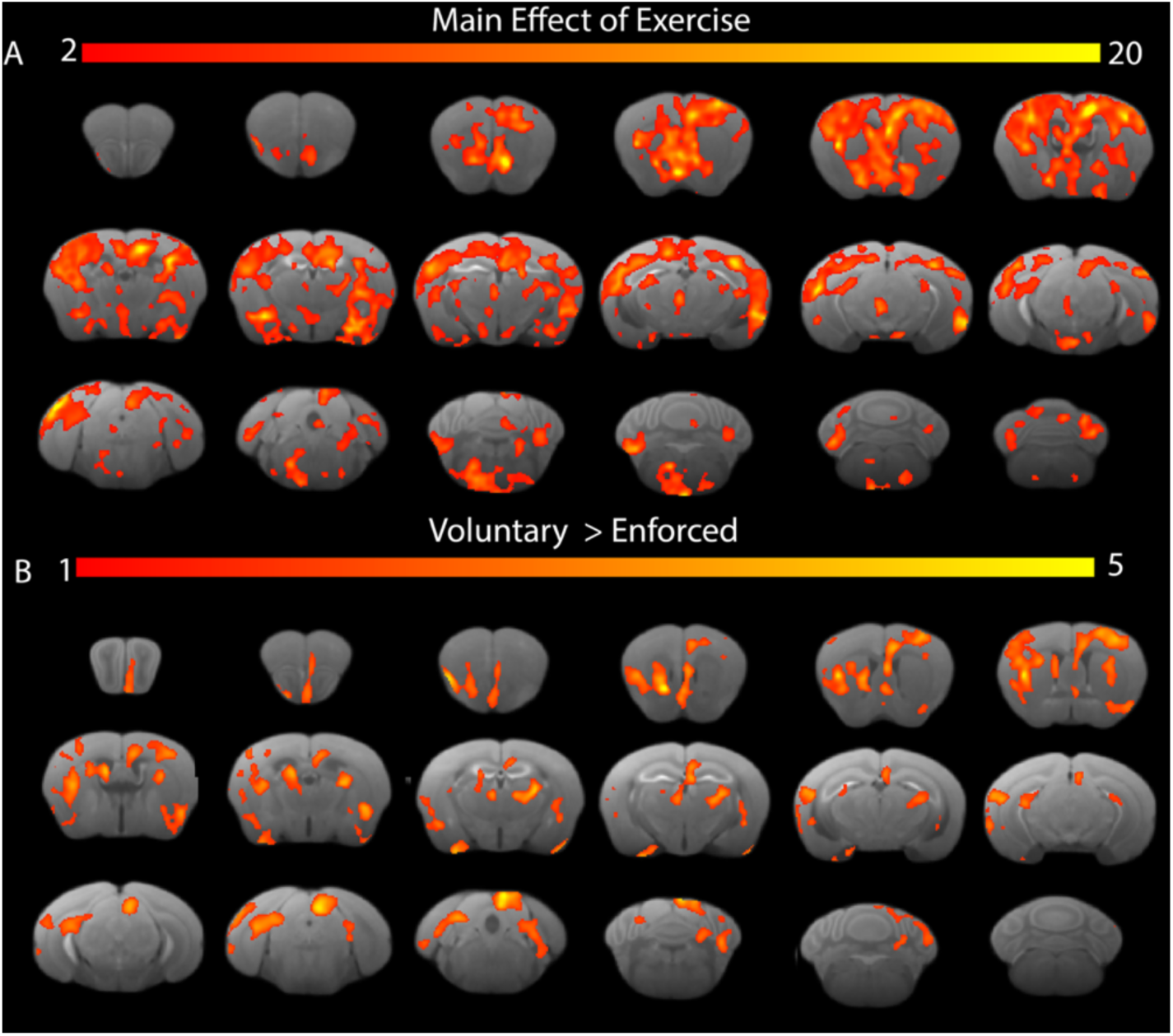
Exercise-associated effects on relative cerebral perfusion. Significant effects of exercise were noted in the olfactory, orbital cortex, M1 and S1, V1, cingulate cortex, and hypothalamus, amygdala, ventral thalamus, superior colliculus, and brainstem reticular nuclei. Voluntary trained mice showed higher perfusion in left M1, and dorsal Hc, bilateral caudate putamen, piriform cortex, V1, and postsubiculum. FDR=5%.

Sparse logistic regression applied to structural connectomes from exercised and sedentary mice identified 10 discriminative subnetworks (**Figure 9**, **Table 1**), yielding an average misclassification rate of 0.27 across five-fold cross-validation. The most predictive for the exercised condition was a network with a large negative weight, centered on the left cingulate cortex (area 24a′ and 24a)— implicated in motivational drive and cognitive control. Other networks contributing to the exercise condition involved the temporal association and entorhinal cortices, anterior thalamic nuclei, basolateral amygdala, primary somatosensory cortex (S1), inferior colliculus, accumbens, and white matter tracts, including the stria terminalis, corpus callosum, and optic tract. In contrast, networks with small positive weights were more predictive of the sedentary condition, involving the bed nucleus of the stria terminalis (BNST), brainstem sensory-motor nuclei, visual relay regions (e.g., lateral and medial geniculate nuclei, optic tract), and the right insular cortex. Notably, regions beyond classical motor circuits were implicated, including structures related to reward processing, cognitive control, and sensory integration. These findings suggest that exercise preferentially engages cortico-limbic and motivational networks, particularly the cingulate cortex, whereas sedentary behavior may rely more heavily on subcortical sensory integration and passive sensory pathways.This suggests that while exercise engages top-down limbic and motivational circuits, sedentary states may reflect increased weighting of subcortical sensory integration or default sensorimotor tone.

**Figure 9.**
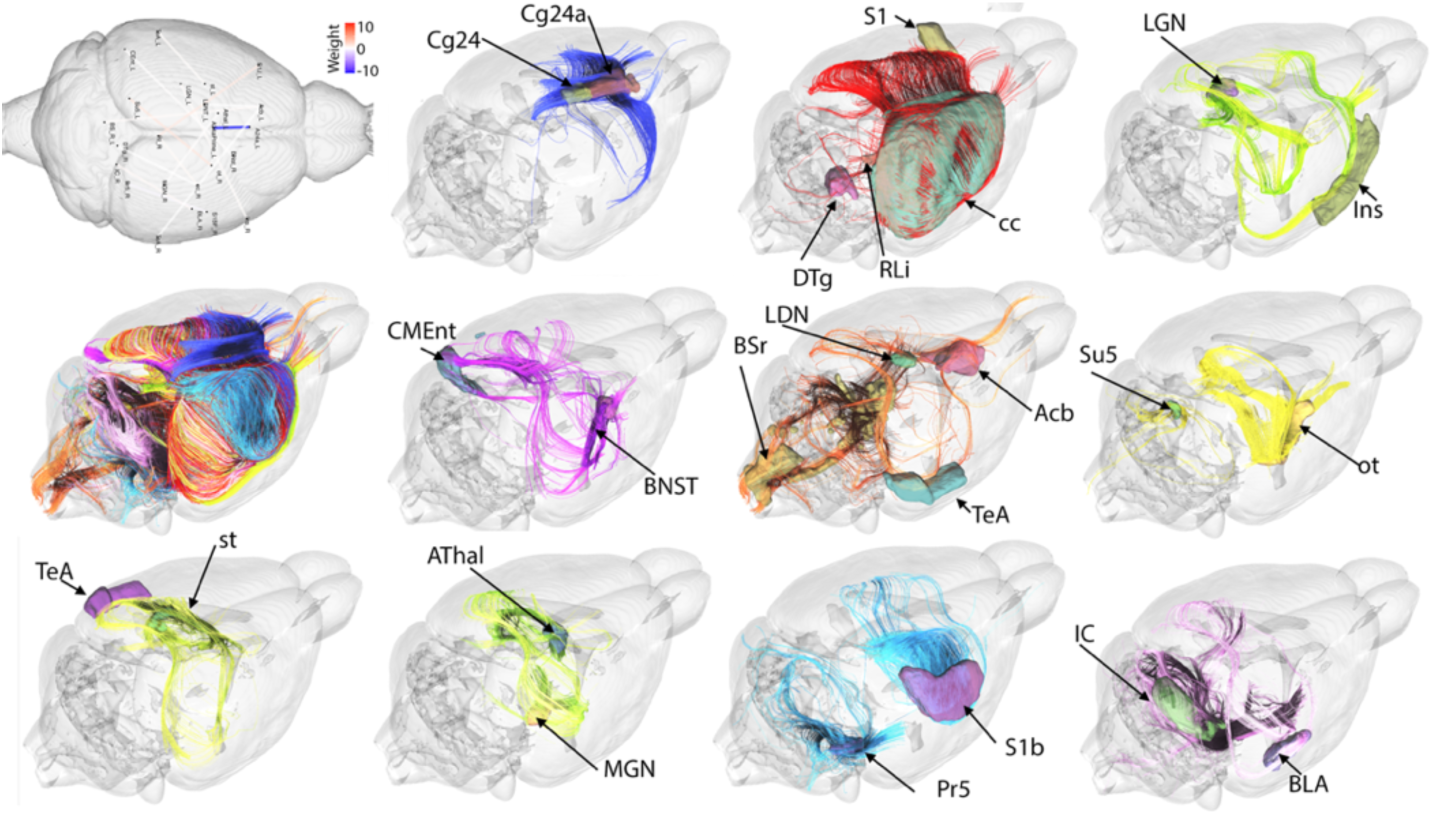
Sparse logistic regression identified networks that discriminated between exercised and sedentary mice. A network centered on the left cingulate cortex area 24a′ and 24a, exhibited the largest negative weight (−10.89), indicating strong association with the exercise condition. Several networks with smaller positive weights were more predictive of the sedentary condition, involving regions such as the bed nucleus of the stria terminalis (BNST), brainstem sensory-motor nuclei, visual relay regions (e.g., lateral and medial geniculate nuclei, optic tracts), temporal association cortex, and the right insular cortex. These results suggest that exercise enhances engagement of cortico-limbic motivational circuits, while sedentary behavior is characterized by greater involvement of subcortical sensory integration pathways. Acb = accumbens; Cg24 = cingulate cortex area 24a; S1 = primary somatosensory cortex; IC = inferior colliculus; BLA = basolateral amygdala; AThal = anterior thalamic nuclei; LGN = lateral geniculate nucleus; MGN = medial geniculate nucleus; BNST = bed nucleus of the stria terminalis; Ins = insula; RLi = rostral linear nucleus; Pr5 = principal sensory nucleus of the trigeminal nerve; Su5 = pedunculotegmental, medial paralemniscal, and supratrigeminal nuclei; ot = optic tract; st = stria terminalis; cc = corpus callosum.

**Table 1.** Brain networks discriminating between exercised and sedentary mice based on structural connectivity.

| Net | Regions | Weight |
| --- | --- | --- |
| 1 | Left Cingulate_Cortex_Area_24a_prime, Left Cingulate_Cortex_Area_24a | -10.89 |
| 2 | Right Rostral_Linear_Nucleus, Right Corpus_Callosum, Left Primary_Somatosensory_Cortex_Jaw_Region, Right Dorsal_Tegmentum | 1.52 |
| 3 | Left Stria_Terminalis, Left Temporal_Association_Cortex | 0.19 |
| 4 | Right Bed_Nucleus_of_the_Stria_Terminalis, Left Caudomedial_Entorhinal_Cortex | 0.06 |
| 5 | Left Latero_Dorsal_Nucleus_of_Thalamus, Left Brain_Stem_Rest, Right Temporal_Association_Cortex, Left Accumbens | 0.25 |
| 6 | Right Insular_Cortex, Left Lateral_Geniculate_Nucleus | 0.17 |
| 7 | Right Optic_Tracts, Left Pedunculotegmental_Medial_Paralemniscal_and_Supratrigeminal_Nuclei | 1.43 |
| 8 | Right Medial_Geniculate_Nucleus, Left Anterior_Thalamic_Nuclei | 0.06 |
| 9 | Right Trigeminal_Sensory_Nucleus, Right Primary_Somatosensory_Cortex_Barrel_Field | -0.36 |
| 10 | Right Basal_Lateral_Amygdala, Right Inferior_Colliculus | 0.32 |

A continuous approach using symmetric bilinear regression (SBL) identified five brain networks associated with the amount of exercise, estimated through the distance run by mice (**Figure 10**). The SBL model significantly outperformed the null model, yielding an RMSE of 294.29 ± 54.42, compared to the null model’s RMSE of 375.03 ± 24.60. Given the standard deviation of running distance (372.87), this result suggests that the identified brain networks effectively predict exercise behavior. The five subnetworks spanned multiple functional domains, including sensory-motor processing, brainstem autonomic control, and limbic circuits related to motivation and memory. Key structures included the medial geniculate nucleus, reticular nucleus, intermediate reticular nucleus, spinal trigeminal nucleus, cuneate nucleus, tegmentum, temporal association cortex, primary somatosensory cortex (S1), pyramidal tracts, inferior cerebellar peduncles, visual cortices (V1, V2), parasubiculum, stria terminalis, septum, and ventral hippocampal commissure. These results suggest that exercise is linked to neuroplasticity across diverse networks, integrating sensorimotor, cognitive, and emotional processes.

**Figure 10.**
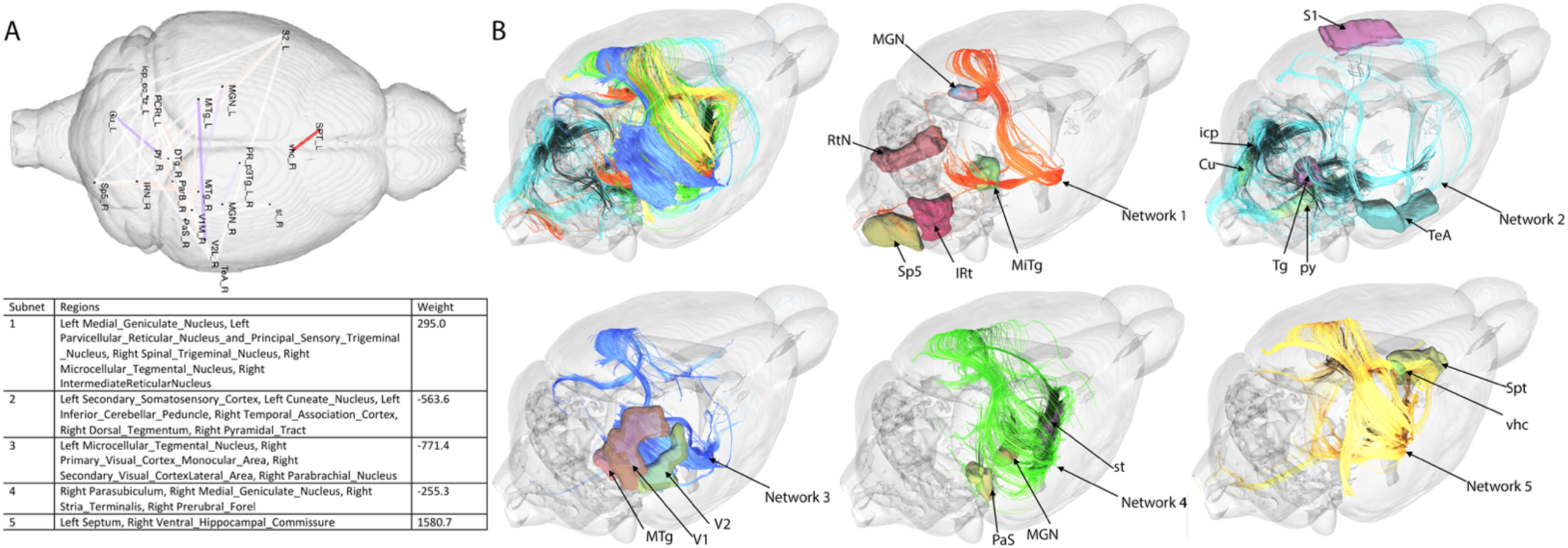
Networks associated with the amount of exercise (i.e. distance run by mice). The symmetric bilinear regression model outperformed the null model with RMSE = 294.29 ± 54.42 while the standard deviation of distance was 372.87, compared to the null model RMSE= 375.03 ±24.6. The five identified subnetworks included the medial geniculate (MGN), reticular nucleus (RtN), intermediate reticular nucleus (Irt), spinal trigeminal nucleus (Sp5); cuneate (Cu), tegmentum (Tg), temporal association cortex (TeA), S1, pyramidal tracts (py), inferior cerebellar peduncles (icp); V1, V2; parasubiculum (PaS), stria terminalis (st); septum (Spt), and ventral hippocampal commissure (vhc).

## Discussion

New evidence supports that lifestyle changes can significantly reduce the risk of developing AD (Livingston et al. 2024). Among these, physical exercise stands out as a key modifiable factor that alters metabolism, brain processes, and cognition, promoting neuroprotection (Baker et al. 2010; Gomez-Pinilla and Hillman 2013; Gomez-Pinilla, Vaynman, and Ying 2008). Exercise may also help mitigate other risk factors, such as obesity and depression. In this study, we show that voluntary and enforced exercise differentially reshape brain structure, function, and behavior in a familial AD mouse model, with notable effects in reducing anhedonia—a core neuropsychiatric symptom of AD. Higher cumulative exercise was associated with greater reductions in anhedonia, suggesting a dose-dependent relationship between physical activity and neuropsychiatric resilience. These findings emphasize the long-term benefits of structured training in sustaining exercise endurance and emotional well-being in a familial AD model.

Multimodal MRI revealed widespread exercise-induced plasticity. In vivo volume increases were observed in motor, entorhinal, and piriform cortices, hippocampus, and cerebellum. Voluntary exercise preferentially enhanced cortical and limbic regions (e.g., cingulate cortex, dentate gyrus), whereas enforced exercise more strongly impacted subcortical and sensory areas (e.g., S1, medial geniculate, and motor regions), as well as the temporal association cortex—a region involved in multisensory integration, semantic memory, and affective processing. These patterns suggest that voluntary exercise promotes cognitive and emotional regulation, while enforced exercise strengthens sensory-motor integration, auditory processing, and autonomic coordination. The in vivo findings underscore the broad neuroprotective effects of exercise, while ex vivo volumetry revealed additional changes in white matter tracts—such as volume increases in the anterior commissure and facial nerve—consistent with structural reinforcement of interhemispheric and cranial motor connectivity. Notably, the cerebral peduncle exhibited reduced volume in vivo, potentially reflecting enhanced efficiency of descending motor output, whereas the anterior commissure showed increased volume ex vivo, suggesting growth of interhemispheric associative fibers. The preferential engagement of cortical networks with voluntary exercise may inform strategies for preserving executive function and memory, while the subcortical and associative enhancements observed with treadmill exercise highlight its potential in rehabilitation for movement disorders, sensory processing deficits, and mood dysregulation, including reductions in anhedonia.

Voluntary exercise induced distinct microstructural adaptations that aligned with volumetric changes. Increases in fractional anisotropy (FA) within memory-related tracts such as the corpus callosum, hippocampal commissure, and cingulum suggest enhanced white matter integrity. In contrast, FA reductions in the cingulate cortex, anterior thalamic nuclei, and amygdala may reflect synaptic remodeling rather than neurodegeneration. Regions showing both volume and FA increases (somatosensory cortex, medial lemniscus, primary auditory cortex) suggest enhanced sensory-motor integration. Conversely, volume increases but FA decreases (entorhinal, cingulate and piriform cortex) suggest synaptic reorganization. These changes were detected in formalin-fixed specimens, ruling out transient physiological effects due to hydration or perfusion. While FA decreases can be interpreted as markers of neurodegeneration, they may also represent beneficial axonal remodeling, gliogenesis, or improved metabolic efficiency, which can increase tissue heterogeneity and alter diffusion. Notably, voluntary exercise elicited greater white matter plasticity than combined regimens, reinforcing its effectiveness in promoting neuroadaptive changes in memory and motivation-related circuits. This effect was especially evident in the corpus callosum, cingulum, fornix, zona incerta, hippocampus, and cingulate cortex. The greater efficacy of voluntary exercise may stem from enhanced engagement of motivation and reward pathways and reduced physiological stress, both of which support neuroadaptive processes. These findings highlight the importance of exercise modality in shaping brain microstructure, with voluntary exercise emerging as a stronger driver of white matter plasticity in networks associated with memory and cognition. These results support the idea that voluntary exercise alone induces more pronounced and functionally relevant microstructural adaptations in white matter, while enforced exercise may impose different, possibly competing mechanisms of plasticity. This aligns with previous research indicating that voluntary exercise is more effective in driving experience-dependent changes in neuroanatomy, potentially due to differences in engagement, motivation, and task-relevant motor learning processes (Baker et al. 2010; Gomez-Pinilla and Hillman 2013; Gomez-Pinilla, Vaynman, and Ying 2008). Future investigations using functional imaging and histological validation will be necessary to clarify the cellular and behavioral consequences of different exercise impacts on microstructure.

T1-weighted MEMRI analyses revealed increased signal in associative and sensory cortices (frontal association, piriform), suggesting enhanced metabolic activity and structural remodeling. In contrast, volume reductions with increased T1w signal in the frontal cortex suggest synaptic pruning, and potentially higher cortical efficiency. In limbic regions (stria terminalis, thalamic reticular nucleus), increased T1w signal, with reduced in vivo volume suggests axonal remodeling and sensory gating refinement, aligning with behavioral improvements. Meanwhile, brainstem and autonomic circuits showed coordinated reductions in both T1w signal and volume, suggesting increased neural efficiency supporting autonomic control, motor regulation, and physiological homeostasis. Increased Mn uptake in the CSF of exercised mice may reflect enhanced neuronal activity, glymphatic clearance, or changes in blood-brain barrier (BBB) permeability. These findings highlight exercise’s ability to modulate brain activity and metal homeostasis, reinforcing its role in cognitive resilience and AD risk mitigation.

It is widely accepted that aging and dementia, together with CVD risk factors such as hypertension and APOE4 carriage, contribute to decreased cerebral perfusion, in particular in vulnerable regions, such as the cortex, thalamus, hippocampus, and white matter in people as well as animal models of AD (Bracko et al. 2021). On the contrary, exercise significantly increased brain perfusion, and memory function reinforcing its neuroprotective role (Tomoto et al. 2023). In our study we found that exercise modulated cerebral blood flow (CBF) in a region-specific manner. Voluntary exercise increased relative CBF in motor, limbic, and associative cortices, including the dorsal hippocampus, cingulate cortex, and piriform cortex—regions also exhibiting structural expansion. Thes results suggest that neurovascular support underpins exercise-driven plasticity, and neuroprotection. Enforced exercise enhanced perfusion in subcortical structures such as the thalamus, ventral hippocampus, as well as auditory cortex. These effects suggest that exercise promotes neurovascular coupling and metabolic support, particularly in circuits associated with cognition and emotional regulation. We did not observe changes in memory, but in anhedonia reduction, and larger studies may be needed to discern such effects.

Connectivity analyses revealed exercise-responsive networks. A negative weight was observed in a network comprising the cingulate cortex (A24a), nucleus accumbens, brainstem, anterior thalamic nuclei, entorhinal cortex, and basolateral amygdala, suggesting these regions were more strongly associated with the sedentary group. A decreased connectivity in the cingulate cortex and nucleus accumbens in exercised mice may indicate reduced reliance on stress-related or reward-seeking behaviors. This may reflect greater efficiency in reward processing, reducing reliance on stress-induced reward-seeking. Similarly, alterations in the basolateral amygdala and anterior thalamic nuclei, involved in emotional regulation and memory processing, suggest that sedentary mice may exhibit differences in affective and cognitive function, particularly linked to memory, reward, and emotional regulation. Conversely, positive network weights, indicating stronger connectivity in exercised mice, were observed in regions including the rostral linear nucleus, corpus callosum, primary somatosensory cortex, dorsal tegmentum, stria terminalis, and entorhinal cortex. Increased connectivity in the corpus callosum and somatosensory cortex suggests enhanced sensorimotor integration and interhemispheric communication, while changes in the stria terminalis and insular cortex indicate potential improvements in stress regulation and interoception. Additionally, enhanced connectivity in the optic tracts, medial geniculate nucleus, and anterior thalamic nuclei suggests that exercise may modulate sensory and cognitive processing. These findings highlight neuroplastic effects of exercise in pathways involved in sensory-motor coordination, emotional regulation, and cognitive resilience, reinforcing its potential as an intervention for mitigating neurodegenerative risk. Our results suggest that exercise engages top-down limbic and motivational circuits, while sedentary states may rely more heavily on subcortical sensory and autonomic pathways, potentially reflecting a less adaptive neural baseline.

Additionally, total running distance correlated with enhanced connectivity in thalamic, brainstem, and cerebellar networks, strengthening the link between exercise intensity and neural plasticity. The involvement of sensory-motor, brainstem, and limbic structures suggests that exercise exerts widespread effects on movement coordination, autonomic regulation, and cognitive-emotional processing. The involvement of primary somatosensory (S1), pyramidal tracts, and cerebellar peduncles support that exercise enhances motor learning and proprioceptive feedback, while that of brainstem nuclei (reticular nucleus, tegmentum, and spinal trigeminal nucleus) suggests engagement of autonomic and reflexive control mechanisms. The participation of limbic structures (stria terminalis, septum, parasubiculum) suggests that exercise modulates motivation, stress resilience, and memory processing. This is consistent with studies showing that physical activity promotes synaptic plasticity, and reduces stress-related hyperactivity (McEwen, Nasca, and Gray 2016; Wu et al. 2024). Additionally, the presence of visual and auditory relay regions (medial geniculate nucleus, V1, V2) suggest that sensory integration plays a role in guiding motor behavior. These findings support that exercise-induced neuroplasticity extends beyond motor pathways to encompass emotional regulation, sensory processing, and cognitive function. Future studies could explore causal mechanisms assessing whether modulating activity in these networks alters running behavior, or whether differences in these networks predict exercise engagement.

Comparing exercise regimens suggests distinct but complementary benefits. Enforced exercise had stronger behavioral effects, reducing anhedonia and enhancing memory, whereas voluntary exercise produced greater neurovascular benefits, increasing cerebral blood flow (CBF) in the motor cortex, hippocampus, and piriform cortex, promoting brain-wide resilience and preserved cortical volume. While enforced exercise may induce stress in animal models, a combined regimen may balance cognitive, emotional, and vascular resilience. Although CBF changes were not always consistent across overlapping regions, structural modifications dominated, suggesting that exercise’s protective effects may arise from cortical and limbic reorganization rather than direct vascular remodeling. These findings support a multi-modal exercise intervention, where voluntary and enforced exercise drive distinct but synergistic neuroprotective effects.

Our findings align with a growing body of literature demonstrating that exercise exerts neuroprotective effects. Consistent with prior studies, we observed that chronic exercise modulated neuroimaging biomarkers associated with AD, including regional brain volume, microstructural integrity, and cerebral blood flow. The preferential effects of voluntary exercise on cortical and limbic regions, such as the hippocampus, piriform cortex, and cingulate cortex, reinforce previous work highlighting the role of exercise in promoting synaptic plasticity, neurogenesis, and vascular resilience in these areas (Pereira et al. 2007) (Morris et al. 2017). Moreover, our results support the established link between exercise and behavioral improvements, particularly in mitigating anhedonia, a symptom that is increasingly recognized as a critical yet understudied component of AD pathology (Lopez et al. 2003) (Hird et al. 2024). The observed reductions in anhedonia, particularly in the enforced exercise group, corroborate findings that exercise modulates reward circuitry and dopaminergic pathways, providing a potential mechanism for its antidepressant effects (Archer, Josefsson, and Lindwall 2014) (Schuch et al. 2014).

Our study also highlights areas of divergence that merit further investigation. While previous research has largely focused on the hippocampus as the primary mediator of exercise-related neuroprotection (Lourenco et al. 2019) (Vaynman, Ying, and Gomez-Pinilla 2004) our findings suggest a distributed effect across sensory, motor, and subcortical networks. The distinct impact of voluntary versus enforced exercise, particularly in modulating white matter microstructure and connectivity, suggests that different exercise paradigms engage unique neuroplasticity mechanisms. Our observation that enforced exercise led to reductions in fractional anisotropy in the cingulate cortex, anterior thalamic nuclei, and amygdala challenges the interpretation of FA as a strict marker of white matter integrity loss. Instead, these changes may reflect synaptic remodeling, increased dendritic complexity, or enhanced gliogenesis. This complexity underscores the need for multimodal imaging approaches to disentangle the diverse mechanisms underlying exercise-induced plasticity. Additionally, our results suggest that the impact of exercise on cerebral blood flow is not uniform across regions or exercise types. While voluntary exercise led to significant increases in CBF in motor and limbic areas, enforced exercise preferentially enhanced perfusion in subcortical structures such as the thalamus and brainstem. This contrasts with prior work that has largely focused on global CBF and suggests that different exercise modalities may differentially engage neurovascular regulatory mechanisms. Understanding such nuances is critical for optimizing exercise interventions in populations where vascular dysfunction is a hallmark of AD progression (Prins and Scheltens 2015).

Many studies have examined the effects of exercise on the brain structure (Biedermann et al. 2016; Draganski and May 2008; Frederiksen et al. 2018; Hayes et al. 2013), both in humans and in rodents (Sumiyoshi et al. 2014); but fewer have examined the impact of different types of exercise on reducing risk for AD. Volume increases in the somatosensory, motor, parietal association, and visual cortices have been reported as soon as after 7 days of training(Sumiyoshi et al. 2014), but no subcortical effects have been reported in that study. Volume reductions have also been reported in regions such as amygdala and hippocampus due to stressors(Ryoke, Hashimoto, and Kawashima 2024), underlining a bidirectional plasticity in adult brains, in regions vulnerable to neurodegeneration. Besides gray matter, white matter changes (Prakash et al. 2010; Voelcker-Rehage and Niemann 2013) may contribute to increased network efficiency.

Understanding the mechanisms underlying training-induced neuroplasticity, and the interactions with genotypes, will inform personalized interventions. An important mechanisms may relate to vascular adaptations to exercise, which have involved the hippocampal dentate gyrus (Clark et al. 2009), which may protect against cognitive decline. For example, exercise-induced upregulation of vascular endothelial growth factor (VEGF) facilitates angiogenesis (Fabel et al. 2003; Lou et al. 2008), supporting neuronal health in the aging brain (Ceci et al. 2024). Another mechanism involves microstructural remodeling in gray matter due to synaptic events and the formation of new connections by dendritic spine growth and change in existing connections (Draganski and May 2008) (Trachtenberg et al. 2002) (Chklovskii, Mel, and Svoboda 2004). A third mechanisms may relate to transport. MEMRI in mouse models can reflect adverse effects of aging on neuronal Ca^2+^ regulation, which are modulated by genetic (mutations in presenilin, α-synuclein, huntingtin, or Cu/Zn-superoxide dismutase; apolipoprotein E), diet, or exposure to toxins (Mattson 2007), all factors which increase risk for AD. Furthermore, these localized changes in brain structure and function may have different dynamics that we need to better understand, to tailor specific interventions that support cognitive resilience. Finally, exercise can mitigate anhedonia, which often accompanies AD, and may be modulated by the structure and activity of reward circuits. Research indicates that regular physical activity may mitigate the effects of the APOE4 genotype (Angelopoulou et al. 2021) on cognitive and vascular function (Kaufman et al. 2021), metabolism (Rhea, Raber, and Banks 2020), and brain volume preservation (Honea et al. 2009), suggesting that exercise could be particularly beneficial for individuals carrying this genetic risk factor, and for females (Palmer et al. 2022).

A limitation of our study is that single-shell diffusion has limited sensitivity for gray matter regions, crossing fibers and complex microarchitecture. Multishell acquisitions, which allow for more accurate signal modeling and improved separation of isotropic and anisotropic diffusion components, would provide a more precise characterization of exercise-driven changes in both gray and white matter. Future studies leveraging multishell aquisitions could enhance our ability to detect microstructural adaptations and the complex interactions between cortical plasticity, connectivity, and behavior. Furthermore, while our study utilized pseudo-continuous arterial spin labeling (pCASL) for cerebral blood flow measurements, it has limited spatial resolution. Improved perfusion sequences with higher signal-to-noise ratios and better motion correction, such as multi-delay ASL or dynamic susceptibility contrast (DSC) MRI, could provide a more detailed assessment of exercise-induced vascular adaptations. These methodological advancements would help clarify how different exercise intensities influence neurovascular function and allow for more precise correlations between perfusion, structural plasticity, and behavioral outcomes. While MEMRI provides enhanced SNR, it also alters T1 relaxation times. To minimize bias in perfusion quantification, we estimated M₀ from ASL control images and interpreted CBF changes in relative rather than absolute terms. This mitigates confounds from manganese-induced T1 shortening, which could otherwise underestimate CBF. Finally, in vivo and ex vivo imaging findings did not always align, highlighting the complexity of exercise-induced neuroplasticity and the importance of using complementary imaging approaches. While in vivo MRI revealed robust volumetric increases in motor, sensory, and associative regions, ex vivo analyses identified more localized effects in hippocampal and subcortical structures, as well as white matter tracts. These discrepancies may reflect differences in hydration status, tissue fixation, and resolution, underscoring the need for integrative analyses when interpreting structural and microstructural brain changes. Taken together, our study reinforces the growing consensus that exercise exerts neuroprotective effects, and highlights the complexity of these adaptations. The differential effects of voluntary and enforced exercise suggest that a one-size-fits-all approach may not be optimal for AD prevention and treatment. Future studies should aim to refine exercise protocols to maximize cognitive and neurovascular benefits while minimizing potential stress-related effects. The observed region-specific effects reinforce the importance of tailored exercise interventions for neurorehabilitation, particularly for aging, neurodegenerative diseases, and cognitive disorders.

## Conclusion

Integrating multimodal imaging with molecular and behavioral analyses offers a framework for elucidating the mechanisms underlying exercise-driven resilience and refining strategies to mitigate AD risk. Our findings highlight distinct yet complementary adaptations in sensory-motor, limbic, and cognitive networks. By advocating for personalized exercise strategies, this work reinforces the potential of lifestyle interventions to enhance cognitive resilience and neurovascular health. Ultimately, exercise emerges as a robust, non-pharmacological neuroprotective strategy. Tailoring regimens to engage cortical versus subcortical circuits—and balancing motivational engagement with physical intensity—may optimize brain health and emotional resilience across aging and neurodegenerative disease trajectories.

## Acknowledgments

We thank Yi Qi, Gary Cofer, Wyatt Austin, and Chris Petty for technical support. We thank Dr Luisa Ciobanu for providing the FISP sequence and helpful discussion.

## Supplementary Tables and Figure Legends

**Supplementary Table 1** The mean values, standard deviations, effect sizes (Cohen’s F), and statistical comparisons for anhedonia reduction, and distances run.

**Supplementary Table 2.** The mean values, standard deviations, effect sizes (Cohen’s F), and statistical comparisons for in vivo volumes, and ex vivo volumes.

**Supplementary Table 3.** The mean values, standard deviations, effect sizes (Cohen’s F), and statistical comparisons for fractional anisotropy.

**Supplementary Table 4.** Mean signal intensities, standard deviations, effect sizes (Cohen’s F), and statistical comparisons for relative T1-weighted MEMRI across brain regions.

**Supplementary Figure 1.** Exercise treatment had a significant effect on body mass (p<0.02), and there was a significant interaction of treatment by time (p<0.0002). At 36 weeks, no significant differences were observed between groups. At 52 weeks voluntary + enforced mice showed significantly lower body mass compared to sedentary controls, while the two exercise groups did not differ significantly. These results suggest that chronic exercise mitigates age-related weight gain in this AD mouse model.

**Supplementary Figure 2.** Main effect of exercise on ex vivo region-based volumetry.

**Supplementary Figure 3.** Main effect of exercise on ex vivo voxel-based volumetry was found in large portions of the isocortex, including M1/S1, hippocampus, subiculum, and cerebellum (A). Voluntary exercised mouse had relatively larger regional volumes in the frontal association cortex, cingulate cortex, M1, ectorhinal and entorhinal cortex, hippocampus, and in particular the dentate gyrus, as well as cerebellum. Mice subjected to enforced exercise had enlarged olfactory areas, M1/S2, parasubiculum.

