## Supplementary figures and images for "Neuroimaging Biomarkers of Neuroprotection: Impact of Voluntary versus Enforced Exercise in Alzheimer’s Disease Models"

### SupplFig1

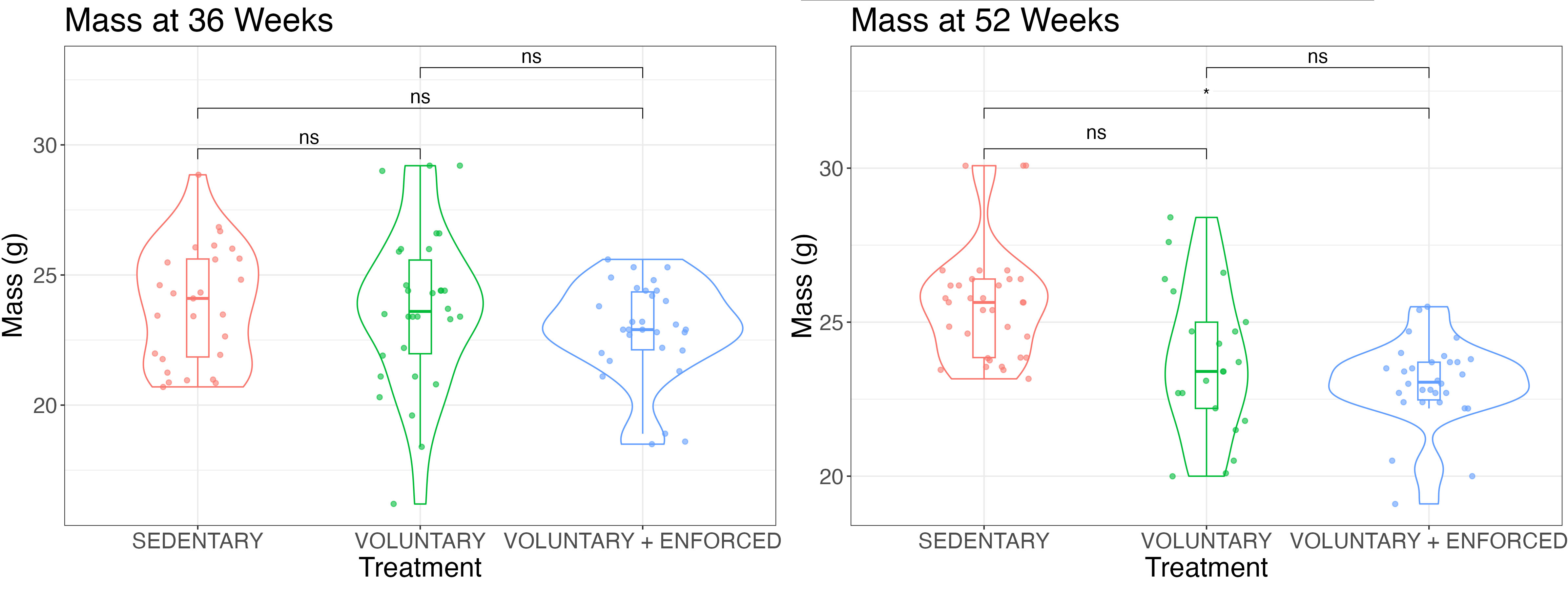

### SupplFig2

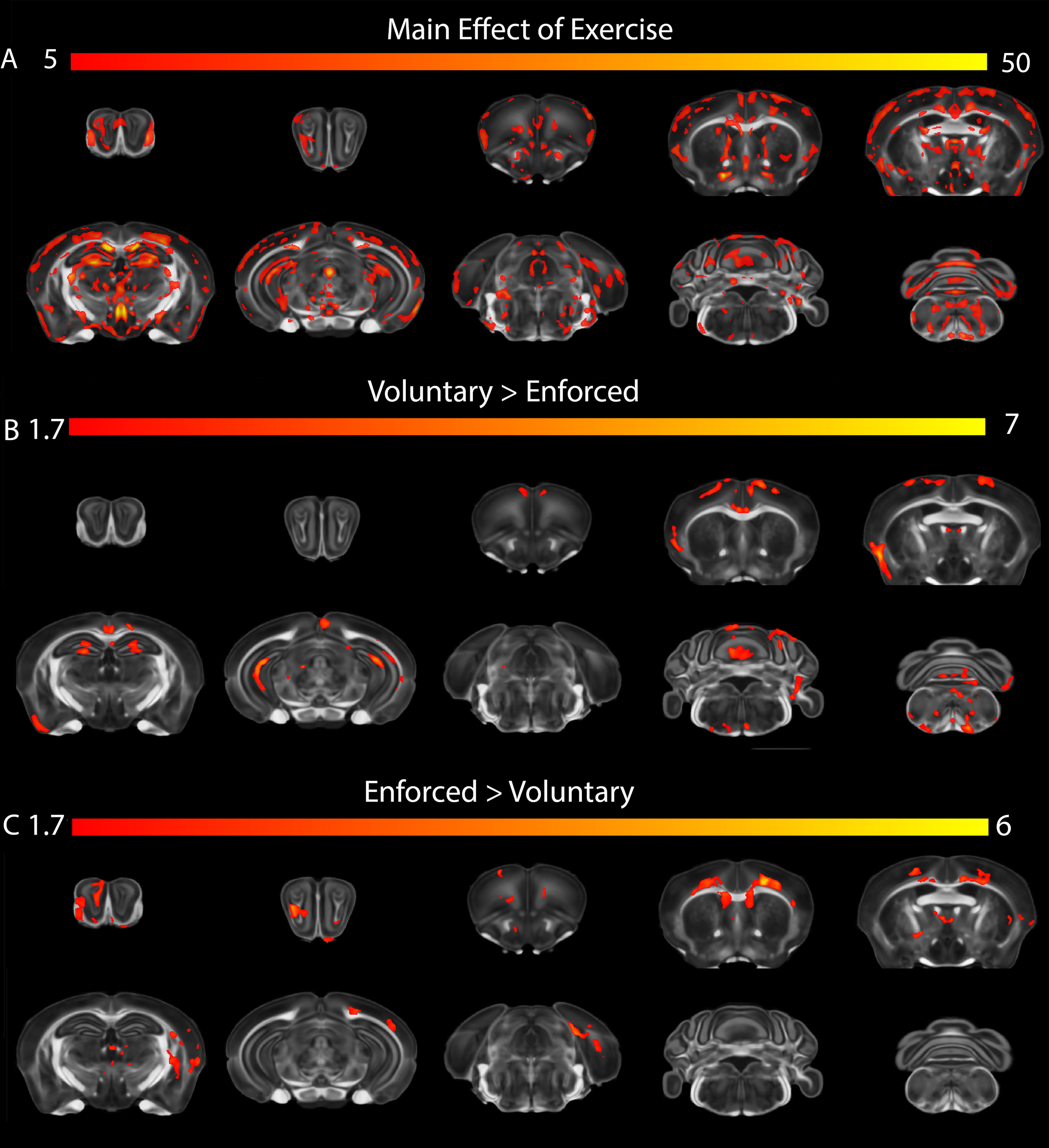

### SupplFig3

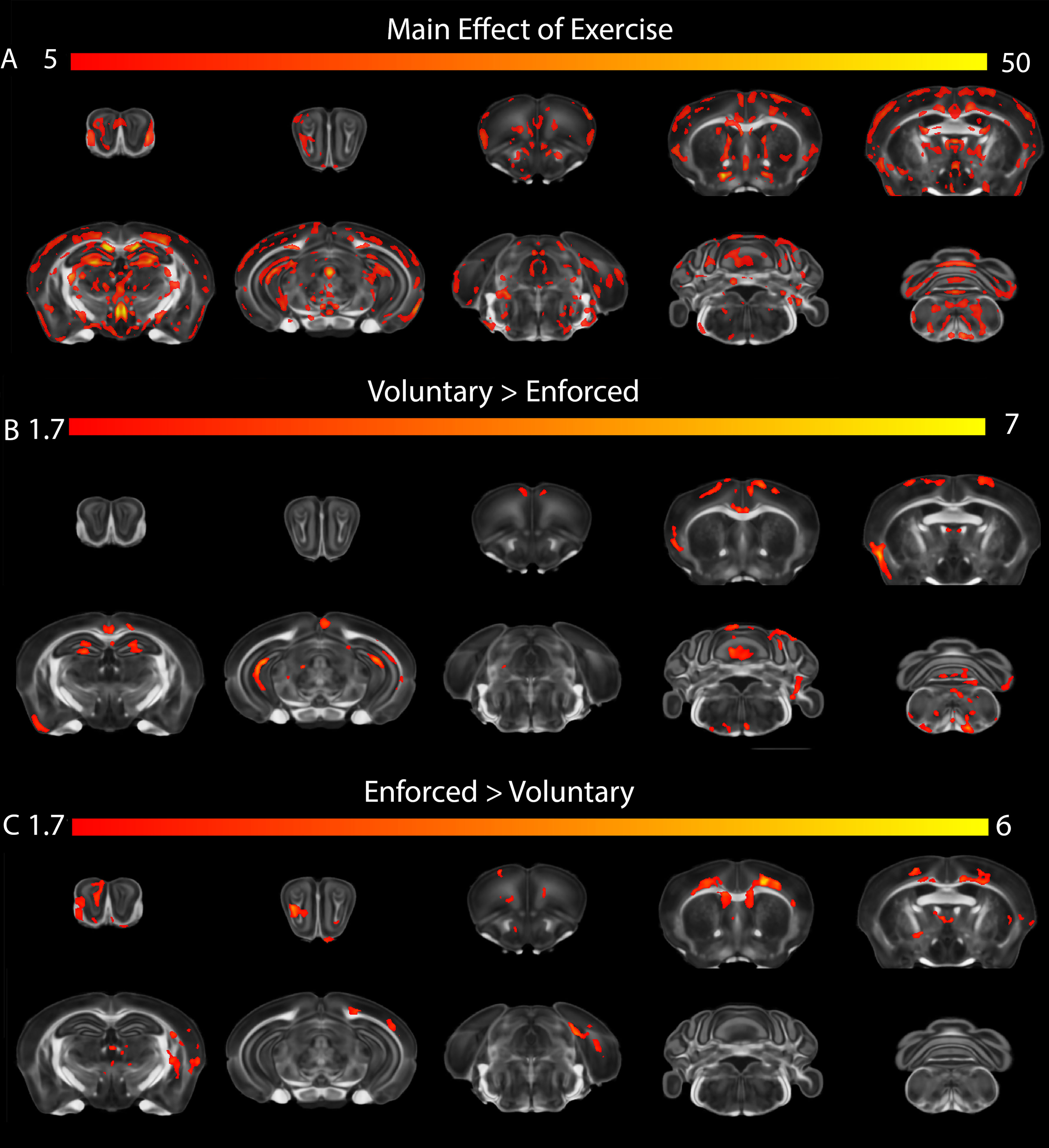
